# Chromosome-level baobab (*Adansonia digitata*) genome illuminates its evolutionary insights

**DOI:** 10.1101/2024.04.14.589434

**Authors:** Justine K. Kitony, Kelly Colt, Bradley W. Abramson, Nolan T. Hartwick, Semar Petrus, Emadeldin H. E. Konozy, Nisa Karimi, Levi Yant, Todd P. Michael

## Abstract

Baobab, *Adansonia digitata*, is a long-lived tree endemic to Africa that holds great economic, ecological, and cultural value. However, our knowledge of its genomic features, evolutionary history, and diversity is limited, rendering it orphaned scientifically. We generated a haploid chromosome-level reference genome anchored into 42 chromosomes for *A. digitata*, as well as draft assemblies for a sibling tree, two trees from distinct locations in Africa, and a related species, *A. za* from Madagascar. Unlike any other plant to date, DNA transposable elements (TEs) make up 33% of the *A. digitata* genome compared to only 10% long terminal repeat retrotransposons (LTR-RTs), which are usually predominant in plant genomes. Baobab has undergone a whole genome duplication (WGD) shared with the Malvoideae ∼30 million years ago (MYA), as well as a confirmed autotetraplody event 3-4 million MYA that coincides with the most recent burst of TE insertions. Resequencing 25 *A. digitata* trees from Africa revealed three subpopulations that suggest gene flow through most of West Africa but separated from East Africa. Gene enrichment analysis for baobab-specific and high fixation index (Fst) suggested baobab may have retained multiple copies of circadian, light and growth genes to coordinate genome protection for longevity through the *UV RESISTANCE LOCUS 8* (*UVR8*) and synchronizing flower development with pollinators. This study lays the groundwork for the creation of breeding resources and the conservation of baobab biodiversity.

## Introduction

The African baobab (*Adansonia digitata*) is a deciduous tree belonging to the Malvaceae family, specifically within the Bombacoideae subfamily. Hereafter, it will be simply referred to as "baobab," with other species like the Australian or Malagasy baobab mentioned as needed in the text. The word “baobab” comes from the Arabic name ‘buhibab,’ which means a fruit with many seeds ^1^. One of the earliest references to baobab was made by Ibn Batuta in the 14th century, who described it as a food and a large, long-living tree in Africa ^2,3^.

Colloquially, baobab is referred to as the ‘upside down tree’ since when it loses its leaves, the branches look like roots; in addition, due to the Hollywood blockbuster “The Lion King,” baobab is also referred to as the “Tree of Life.”

Baobab offers various edible parts, i.e., seeds, leaves, roots, flowers, and powdery fruit pulp. The fruit is particularly rich in vitamin C, antioxidants, anti-inflammatory compounds, minerals, and fiber. Beyond its dietary benefits, the bark is used in crafting robes and mats, adding to the economic importance of the baobab tree. Furthermore, the seeds yield oil used in cosmetics ^4^. Baobab seeds contain phytic acids, just like legume seeds ^5^; however, proper processing can reduce these acids ^6^. The recent approval of the powdery fruit pulp as a food ingredient by the European Commission (EC) and the United States Food and Drug Administration (FDA) has significantly increased demand for baobab products outside of Africa. The estimated value of the baobab powder market worldwide was US$8.2 billion in 2022, and is anticipated to expand to US$12.1 billion by 2030 ^7^. Thus, there is economic interest and social need for genomic resources to *study*, *preserve*, and *increase* baobab yields ^8^.

Baobabs are some of the oldest and largest non-clonal living organisms on Earth with trees that can live over 2,400 years with canopy sizes of greater than 500 m^3^ and trunks reaching diameters of up to 10.8 meters (35 feet) ^9^. However, baobabs are unlike most large and long-lived trees; they are succulents characterized by parenchyma-rich tissues that efficiently store water and therefore, do not form “growth rings” or true wood ^10^. Achieving maturity in the wild presents a considerable challenge for baobab trees since seedlings face predation from caterpillars, goats, and cattle ^11^. Despite having bisexual flowers, baobabs are mostly self-incompatible, depending on external pollinators for successful fertilization ^12,13^. In natural populations, *A digitata* is primarily pollinated by bats ^14,15^, with occasional visits by bushbabies ^16^, and hawkmoths in southern Africa ^13^. Since *A. digitata* is an obligate outcrosser, the populations harbor a high level of diversity, which is observed as heterozygosity in the genome ^13,17^. The baobab tree typically exhibits slow growth to maturation, requiring anywhere from 8 to 23 years to reach the flowering stage, which can impede initiatives aimed at pre-breeding and genetic characterization ^18,19^.

Despite the baobab tree’s remarkable ability to thrive in harsh environments with less than 500 mm of annual rainfall and sandy, rocky soils, recent reports suggest elevated mortality in the oldest trees ^9,20^. A 1,400 year old tree called Chapman baobab in the Makgadikgadi Pans (MP) National Park of Botswana, thought to be the cradle of *Homo sapiens* ^21^, suddenly died on January 7, 2016 along with eight other historic baobabs, including the oldest known baobab, Panke ^9^. Climate change in southern Africa is implicated in the sudden deaths of these giants ^22^, yet the specific drivers remain a mystery. Numerous hypotheses have emerged regarding the causes of these deaths, attributing them to rising global temperatures, pathogen infections, soil compaction due to farming activities, or overexploitation ^9,11,20^. However, the lack of comprehensive scientific studies has hindered a conclusive understanding of this phenomenon. A major impediment is the scarcity of high-quality genome resources. A recently released draft baobab genome only reached contig level and fell short of the necessary depth for a comprehensive genetic analysis ^9,18,23,24^.

The *Adansonia* genus has eight recognized species, with six species endemic to Madagascar: *A. grandidieri*, *A. perrieri, A. rubrostipa*, *A. madagascariensis*, *A. za*, and *A. suarezensis*, while *A. digitata* and *A. gregorii* are indigenous to mainland Africa and Australia, respectively ^3,17,20^. While seven baobab species are widely acknowledged as diploids (2n=2x= 88) ^1,17,25^, *A. digitata* is a tetraploid ^26^. Pettigrew et al. (2012) claimed to have found a second diploid species in Africa, *A. kilima*, but this hypothesis has not been supported by subsequent research ^17,27^. Historical reports for African baobab suggested chromosome numbers of 2n=96-168, accompanied by genome sizes ranging from 3 to 7 pg 2C/holoploid ^1,17,28^. However, the ploidy of *A. digitata* remains uncertain, with questions about whether it is diploid, autotetraploid, or allotetraploid ^20^.

Here, we report genome size estimates for all eight recognized baobab species using a K-mer-based method from short-read genomic sequences. This method can provide independent estimates to those obtained previously using Feulgen staining and flow cytometry ^1^. We also generated a haploid chromosome-scale assembly of *A. digitata* (Ad77271a; originally from Tanzania) as well as long read draft assemblies of an Ad77271a sibling (Ad77271b), and offspring of the following trees; the Kord Bao Sudan (AdKB), the Okahao Heritage Tree from Namibia (AdOHT), and an additional species, *A. za* (Aza135) from Madagascar. Finally, we resequenced 25 additional *A. digitata* trees representing different regions of Africa with short reads to assess genetic diversity. These geographically isolated populations highlight genomic variations and play a crucial role in validating the occurrence of baobab polyploidy. The findings from this work deepen our understanding of baobabs and will facilitate future breeding and conservation initiatives.

## Results

### Adansonia species genome sizes and heterozygosity

Feulgen staining and flow cytometry have been used historically to estimate genome size ^29^. Using cytological methods, it was suggested that *A. digitata* has 42 chromosomes with a haploid genome size of approximately 920 megabases (Mb) and a 2C-DNA value of 3.8 pg Here, we skim sequenced all eight recognized baobab species and employed K-mer frequency analysis to estimate the genome sizes of *A. digitata, A. madagascariensis, A. perrieri, A. za, A. gregorii, A. grandidieri, A. rubrostipa,* and *A. suarezensis*. Notably, the genome sizes ranged from 646 megabytes (Mb) in *A. perrieri* to 1.5 gigabytes (Gb) in *A. grandidieri*. The K-mer based genome size estimates were consistent with the previous estimates and suggested variability in genome size among *Adansonia* species ^1,24^. The repeat content from the K-mer estimates varied between 25.8% and 57%, with *A. rubrostipa* having the highest repeat content. Additionally, heterozygosity levels ranged from 1.1% to 1.9%, with *A. digitata*, which is distributed across different ecologies in mainland Africa, exhibiting the highest heterozygosity of 1.9%. This high heterozygosity level could be attributed to *A. digitata* outcrossing ^13,17^ and possible autotetraploidy ^26^ (Table 1).

**Table 1:** K-mer based genome size estimates across the eight *Adansonia* species.

|  | *A. digitata* | *A. madagascariensis* | *A. perrieri* | *A. za* | *A. gregorii* | *A. grandidieri* | *A. rubrostipa* | *A. suarezensis* |
| --- | --- | --- | --- | --- | --- | --- | --- | --- |
| genome size (bp) | 749,665,340 | 647,683,220 | 645,677,030 | 647,031,624 | 701,880,489 | 1,528,554,306 | 1,336,192,477 | 752,433,438 |
| Unique genome (bp) | 528,590,582 | 480,642,287 | 470,932,522 | 471,700,912 | 507,860,224 | 1,043,221,753 | 574,147,066 | 526,812,479 |
| repeat genome (bp) | 221,074,757 | 167,040,933 | 174,744,508 | 175,330,713 | 194,020,266 | 485,332,552 | 762,045,411 | 225,620,959 |
| repeat fraction (%) | 29.5 | 25.8 | 27.1 | 27.1 | 27.6 | 31.8 | 57 | 30 |
| Heterozygosity (%) | 1.9 | 1.5 | 1.6 | 1.6 | 1.2 | 1.1 | 1.8 | 1.1 |

### Baobab genome assembly, annotation and ploidy estimation

The Germplasm Resources Information Network (GRIN) is responsible for the conservation of an *A. digitata* tree named "PI77271" (Supplementary Fig. 1). We chose PI77271 to sequence since it will be available to the scientific community for further experiments. We long read Oxford Nanopore Technologies (ONT) sequenced two PI77271 sibling seedlings, which we named "Ad77271a" and "Ad77271b," to examine heterozygosity and explore possible polyploidy in *A. digitata*. The genome assembly of Ad77271a and Ad77271b resulted in 1,780 and 2,430 contigs, with N50 lengths of 15.3 and 15.0 Mb respectively (Table 2). Ad77271a was scaffolded into 42 chromosomes using High Throughput Chromatin Conformation and Capture (HiC) (Fig. 1c), aligning with the known number of haploid chromosomes (2n = 4x = 168) in *A. digitata* ^1,17^. The final haploid genome sizes were 674 Mbp (Ad77271a) and 678 Mbp (Ad77271b) (Table 2), congruent to a recently published *A. digitata* genome assembly (686 Mb) based on short reads ^24^.

**Fig. 1:**
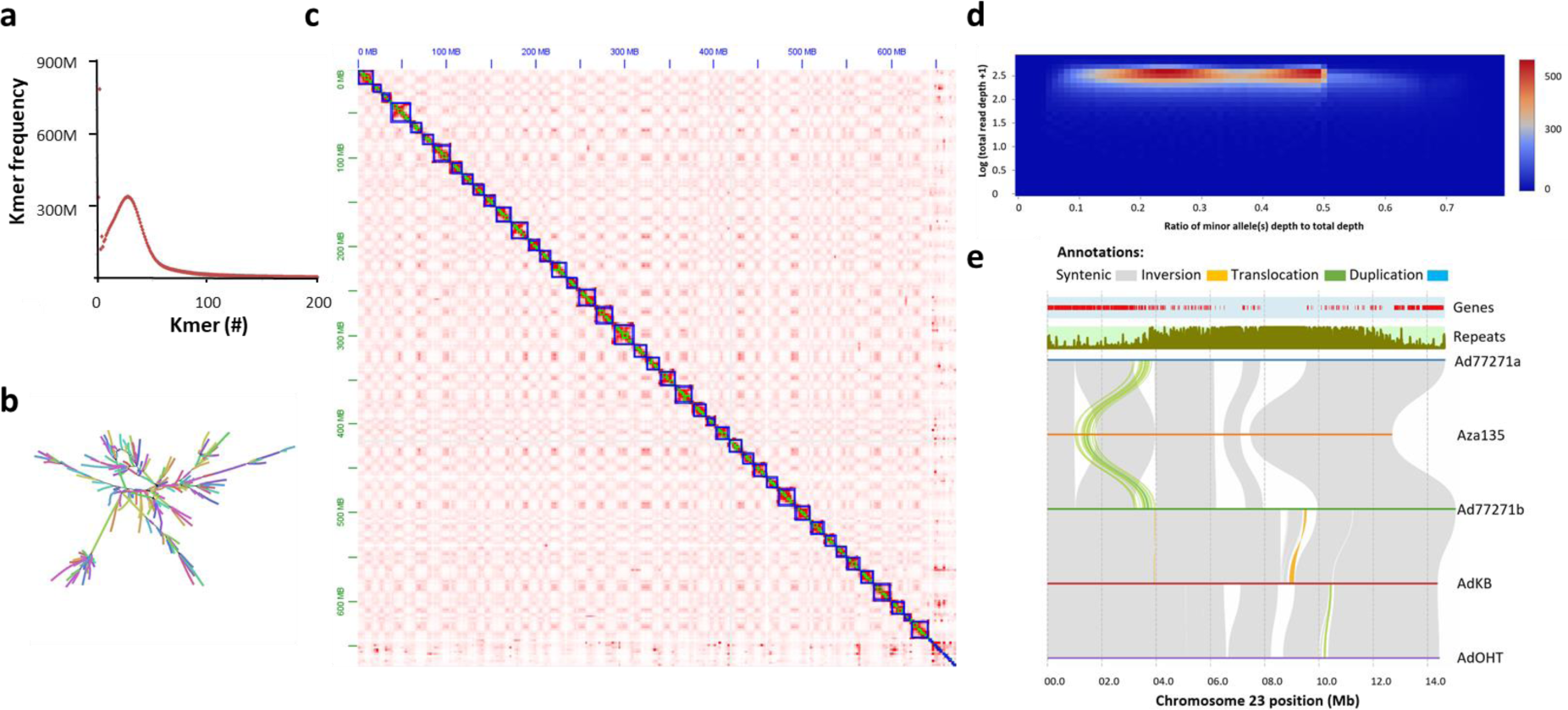
Characteristics of baobab (*Adansonia digitata*) genome. **a** K-mer frequency analysis. A distinctive unimodal pattern suggests a diploid homozygous genome with a size of 750 Mb. **b** Assembly graph of Ad77271a suggests that there are both stretches of heterozygosity as well as transposable elements (TEs) that impact genome assembly. **c** HiC contact map of *A. digitata* (Ad77271a) shows the 42 chromosomes and the shared centromere sequence across the chromosomes. The bottom right corner was unscaffolded centromere sequence. **d** Two-dimensional histogram depicts tetraploid based on minor allele frequency coverage. For diploid organisms, a single peak is expected. However, for tetraploid organisms, the histogram should exhibit two peaks, approximately located at 0.25 and 0.5, respectively. **e** Structural rearrangements and synteny between *A. digitata* (Ad77271a, Ad77271b, AdKB, and AdOHT) and *A. za* (Aza135). A translocation on chromosome 23 distinguishes *A. digitata* from *A. za* species. Gray, orange, green, and blue ribbon colors represent syntenic, inversion, translocation, and duplication structural variations, respectively. The tracks above the structural variant ribbons in the panel depict the distribution of genes and repeat sequence along chromosome 23.

**Table 2:** Statistics for four *Adansonia digitata* genome assemblies (Ad77271a, Ad77271b, AdKB and AdOHT) and *Adansonia za* (Aza135)

| Name | Ad77271a | Ad77271b | Aza135 | AdKB | AdOHT |
| --- | --- | --- | --- | --- | --- |
| Total length (bp) | 674,036,095 | 677,910,090 | 600,003,679 | 682,236,562 | 704,441,320 |
| Contigs (#) | 1,780 | 2,430 | 7,173 | 7,348 | 4,783 |
| Total GC content (%) | 32 | 32 | 31 | 32 | 31 |
| Mean length(bp) | 379,247 | 279,392 | 83,648 | 92,847 | 147,280 |
| N50 (bp) | 15,293,049 | 14,940,500 | 703,193 | 821,818 | 631,186 |
| Copia (#) | 29,501 | 26,820 | 21,861 | 33,438 | 25,453 |
| Gypsy (#) | 23,207 | 41,275 | 17,855 | 25,794 | 10,618 |
| LTR_unknown (#) | 35,805 | 36,424 | 16,450 | 42,141 | 58,288 |
| CACTA (#) | 39,269 | 42,581 | 39,263 | 38,336 | 112,016 |
| Mutator (#) | 104,798 | 94,707 | 106,663 | 124,552 | 123,953 |
| PIF_Harbinger (#) | 30,821 | 28,021 | 23,360 | 28,328 | 34,576 |
| Tc1_Mariner (#) | 5,997 | 6,455 | 3,858 | 6,290 | 7,431 |
| hAT (#) | 23,225 | 13,545 | 8,152 | 19,246 | 11,542 |
| Helitron (#) | 86,011 | 93,088 | 67,243 | 84,842 | 90,424 |
| Total TE (#) | 378,634 | 382,916 | 304,705 | 402,967 | 474,301 |
| Masked DNA TE (bp) | 222,280,883 | 200,555,550 | 159,891,819 | 234,252,229 | 220,359,705 |
| Masked DNA TE (%) | 33 | 30 | 27 | 34 | 31 |
| Masked LTR-RT TE (bp) | 73,645,538 | 100,143,935 | 51,909,405 | 72,289,272 | 94,452,961 |
| Masked LTR-RT TE (%) | 10 | 14 | 9 | 11 | 13 |
| Total TE (bp) | 295,926,421 | 300,699,485 | 211,801,224 | 306,541,501 | 314,812,666 |
| TE masked (%) | 44 | 44 | 35 | 45 | 45 |
| Genes (#) | 43,983 | 43,490 | 46,890 | 45,412 | 46,734 |
| Mean gene length (bp) | 2,107 | 2,112 | 2,295 | 2,064 | 1,964 |
| Average exon/gene (#) | 4 | 4 | 5 | 4 | 4 |
| BUSCO complete (%) | 98 | 98 | 93 | 98 | 94 |

Utilizing ONT long read sequencing we assembled draft genomes of two additional *A. digitata* trees: Kord Bao (AdKB) from Sudan and Okahao Heritage Tree (AdOHT) from Namibia. Furthermore, we assembled the genome of another *Adansonia* species, *A. za* (Aza135) from Madagascar, 23°12’44.0"S 44°02’32.2"E) (Table 2; Supplementary Table 1). The additional assemblies had modest N50 lengths between 600-800 kb as compared to the Ad77271a and Ad77271b genomes (Table 2), but the contiguity was more than sufficient to describe structural variation (SV), nucleotide diversity, and polyploidy. We evaluated the completeness of the assemblies using Benchmarking Universal Single-Copy Orthologs (BUSCO; eudicots-odb10), revealing ∼98% completeness for Ad77271a, Ad7721b, AdKB, and slightly lower completeness for both AdOHT (94%) and Aza135 (93%) (Supplementary Fig. 3a).

Several chromosome rearrangements were found between *Adansonia* species, while the predominant difference between the genomes were due to a lack of alignment in what may be the centromere region (Fig. 1e, Supplementary Fig. 4; see centromere section). The unaligned regions could be due to unassembled repeat sequence, which is common in most genome assemblies still, or it could reflect true differences between the centromeres, specifically between *A. digitata* and *A. za*. Inversion count varied across comparisons, ranging from 28 inversions between Ad77271a and Aza135 to 135 inversions between Ad77271b and AdKB. The comparison between Ad77271b and AdKB showed the highest number of translocations with 359 occurrences, followed closely by AdKB versus AdOHT which had 333 occurrences (Supplementary Table 8). The structural differences were observed despite the overall nucleotides similarity; this has been noted in other autotetraploid genomes ^30^.

The only large (∼1 Mb) translocation identified was between the *A. digitata* species and Aza135 on chromosome 23 (Chr23) that moved ∼120 genes away from the putative centromere region closer to the telomere (Fig. 1e). Gene ontology (GO) enrichment was associated with RNA splicing and the region also included *HISTONE-FOLD COMPLEX 2* (*MHF2*) and *REPLICATION PROTEIN A 1B* (*RPA1B*), which are involved with DNA replication, repair, recombination, and transcription, as well as the later being involved with sustaining telomeric DNA length ^31–34^. In *Arabidopsis thaliana*, loss of the *RPA1B* ortholog is sensitive to Ultraviolet B (UV-B) light with reduced chlorophyll A and B and inhibited root growth, which is due to elevated DNA damage ^33^. It is thought that long lived organisms actively protect their telomere length ^35,36^, and repair their DNA ^37,38^.

Since the K-mer frequency analysis yielded a haploid genome size of approximately 750 Mb (Table 1) and a graph structure indicative of high heterozygosity (Fig.s 1a; b), yet only one peak consistent with a diploid genome, it was important to test the tetraploid designation of *A. digitata* ^1^. The single peak from K-mer analysis may be the result of inter-homeolog recombination as have been noted in other genomes, such as the coast redwood ^39^ and duckweed *Lemna minor* ^40^. We assessed genetic variations within the baobab sibling’s genomes and tested whether *A. digitata* is diploid or polyploid. This comparison unveiled 14.6 million SNPs in Ad77271a against Ad77271b (Supplementary Fig. 5). A heatmap of coverage at heterozygous loci showed two dense peaks at about 0.25 and 0.5, which was what would be expected if it were an autotetraploid organism (Fig. 1d; Supplementary Fig. 5). Due to the similarities between the four subgenomes, additional methods will be required to complete a fully haplotype resolved autotetraploid assembly such as has been done for potato ^30^.

### High DNA transposons accumulation in baobab genome

We conducted an *ab initio* TE prediction in the Ad77271a genome, which resulted in 378,634 transposable elements with a total length of 296 Mb (∼43% of the genome size) (Fig. 2a; Table 2). In most plant genomes, long terminal repeat retrotransposons (LTR-RTs) comprise the largest TEs fraction due to their copy-and-paste mechanism of proliferation that results in genome bloating ^41^. However, in the Ad77271a genome, LTR-RTs only comprised 10% of the TEs complement, while DNA TEs, which proliferate by a cut-and-paste mechanism, made up 33% of the genome (Table 2). Within DNA TEs, ’Mutator-type’ elements were predominant: 115,882,201 (52.1%), while CACTA, PIF Harbinger, Tc1 Mariner, hAT, and helitron elements comprised 26,790,652 (12.1%), 16,680,965 (7.5%), 2,970,466 (1.3%), 14,266,363 (6.4%), and 45,690,236 (20.6%), respectively; this pattern was observed in all the baobab genomes (Fig. 2a; Supplementary Table 2). Moreover, both Copia LTR and CACTA elements showed signs of recent bursts. Investigation of TE insertion times using synonymous substitution (Ks) values showed that most TEs were inserted earlier than 10 million years ago (MYA) based on the peak around 10 MYA. We also found more recent TEs insertions, especially in the autotetraploid *A.digitata*, with a peak around 2-3 MYA (Fig. 2b; Supplementary Fig. 7).

**Fig. 2:**
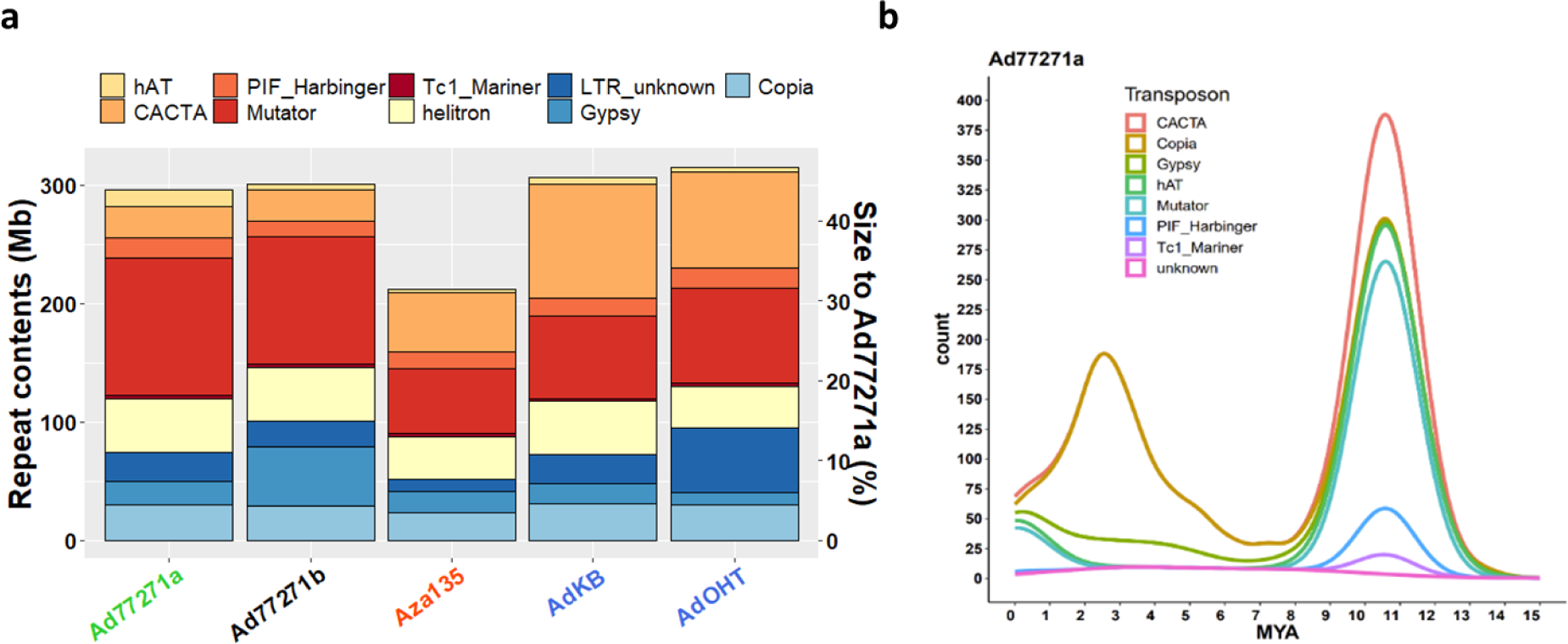
Classification and evolution of transposable elements (TEs) in baobab. **a** Barplot comparison of TEs classification and sizes in five baobab genomes: Ad77271a, Ad77271b, Aza135, AdKB, and AdOHT. Aza135 is highlighted in the center to emphasize TE divergence. Except for Aza135 (*A. za*), the other four genomes belong to *A. digitata*. Distinct colors in the legend denote TE classes. The TE proportion to genome size (Mb) and percentage are displayed on the left and right y-axes, respectively. **b** Density plots of intact TEs, displaying the distribution of different types of TEs. Around 3 - 4 Million Years Ago (MYA), the *Adansonia digitata* genome experienced elevated levels of COPIA long terminal repeat retrotransposons (LTR-RTs).

*Baobab genome organization: centromere, telomere, rDNA and DNA methylation* The 42 chromosomes of baobab were small, ranging from 9 to 23 Mb. The HiC connection map suggested the centromere sequence was highly conserved across the chromosomes, and consistent with this, we found a putative centromere repeat with a base unit of 158 bp, and a higher order repeats (HORs) of 314 and 468 bp (Fig.s 3a; d; e). In general, the centromere arrays assembled well into 1-2 Mb regions that were both metacentric as well as acrocentric (Chr12, 24, 26, 28, 35, 38, 39 are acrocentric; Supplementary Fig. 9). Unlike the allotetraploid *Eragrostis tef*, which has distinct centromere repeats per sub-genome ^42^, we identified one centromere repeat in the *A. digitata* genome consistent with autotetraploidy. In addition, telomere sequences were assembled on the ends of most chromosomes, and these telomeres were long compared to other plants with maximum sequences spanning 30 kb as compared to the model plants *Arabidopsis thaliana* and *Zea mays* that have 3-5 kb telomeres (Fig. 3b; Supplementary Table 3) ^35^. In the Ad77271a genome, we assembled a total of 300 Kb of the ribosomal DNA (rDNA) with only one 26S array that included 35 copies on Chr38, and one 5S array that included 442 copies on Ad77271a Chr01 (Supplementary Table 4).

**Fig. 3:**
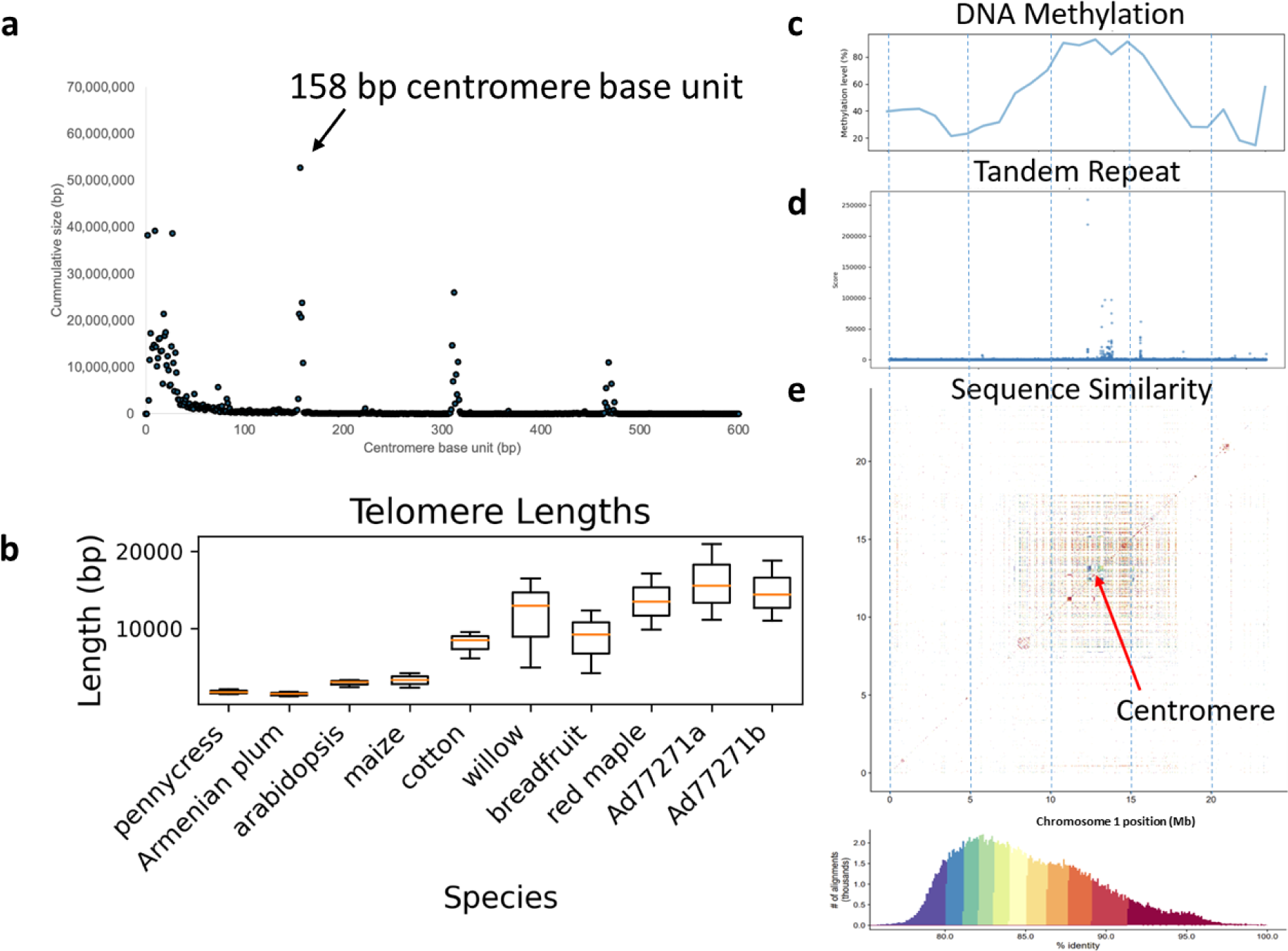
Genomic organization of baobab (*Adansonia digitata*). **a** Predominant and large centromere array with a base unit of 158 bp with higher order repeats (HORs) of 314 and 468 bp. **b** Comparative box plots showing telomere length distribution between baobab (Ad77271a and Ad77271b) and seven other randomly selected plant species. Baobabs have long telomeres (AAACCCT), reaching a maximum length of 30 kb, which aligns with the renowned longevity of baobabs ^23^. **c** Plot showing elevated DNA methylation (5- Methylcytosine) levels in centromeric regions of chromosome 1. **d** Increased occurrence of tandem repeats within the centromeric region. **e** The meta-centromeric region of chromosome 1 is shown on a pairwise sequence identity heatmap. The scale below shows the percentages of sequence similarity. The blue dotted lines show regions along the chromosome.

A consistent marker of age and longevity in animals is DNA methylation ^43,44^. In plants, the situation is more complex but it has been shown that DNA methylation is linked to genome stability and viability ^45^. DNA methylation is sometimes referred to as the fifth base because it is a chemical modification to the cytosine base of DNA that is known in plants to specify cell fate, silence transposable elements, and mark environmental interactions ^46^. Long ONT reads enable the direct detection of 5-methylcytosine (5-mC) DNA methylation ^47^. We leveraged our long ONT reads to look globally at DNA methylation and found average levels of 54.74% in Ad77271a, 54.94% in Ad77271b, and a higher level of 62.52% in AdKB (Supplementary Fig. 8). Consistent with previous findings, we found that there was an increase in DNA methylation in the putative centromere arrays (Fig.s 3c; d; e). Additionally, we found that TEs exhibit hypermethylation, while genes display hypomethylation (Supplementary Fig. 6). These observations align with expected methylation patterns seen in other angiosperms ^48^.

### Gene expansion, contraction and comparative orthogroup analysis

Baobab is unlike most large, long-lived trees because it is succulent, and when it dies its “wood” seems to “deflate,” or mush; wood soaked in water will completely disintegrate after several days, leaving only fibers that are used as packing materials ^10^. We predicted and annotated genes across the baobab genome assemblies to compare against other tree genomes and closely related species like cotton, cacao and bombax. Leveraging both *ab initio* and protein homology gene prediction, we estimated an average of 44,000 genes in *A. digitata* (Ad77271a, Ad7721b, AdKB, and AdOHT), and slightly higher number in Aza135 (46,890).

Gene family comparisons between baobab and fifteen plants identified a total of 40,312 orthogroups (OG) (Fig. 4a). In the Malvaceae, including baobab, cotton, cacao, durio, and cotton tree, alongside representatives from various plant families, identified 3,851 expanded gene families shared between baobab and cacao, 799 gene families expanded in baobab only, and 232 gene families expanded in cacao only (Fig. 4d). Among contracted gene families, 596 were shared between baobab and cacao, with 1,465 specific to baobab and 5,678 specific to cacao (Fig. 4d). While there were 107 and 283, Ad77271a and Ad77271b specific OG respectively, the largest shared specific OGs (4,626) were baobab-specific (both Ad77271a and Ad77271b), suggesting baobab has some unique gene content compared to the other plants chosen for this analysis (Fig. 4a; Supplementary Table 6). The second largest shared group of OG was between monocot genomes, while cotton genomes were the third largest shared OG group with 1,723 and 1,263 respectively (Fig. 4a).

**Fig. 4:**
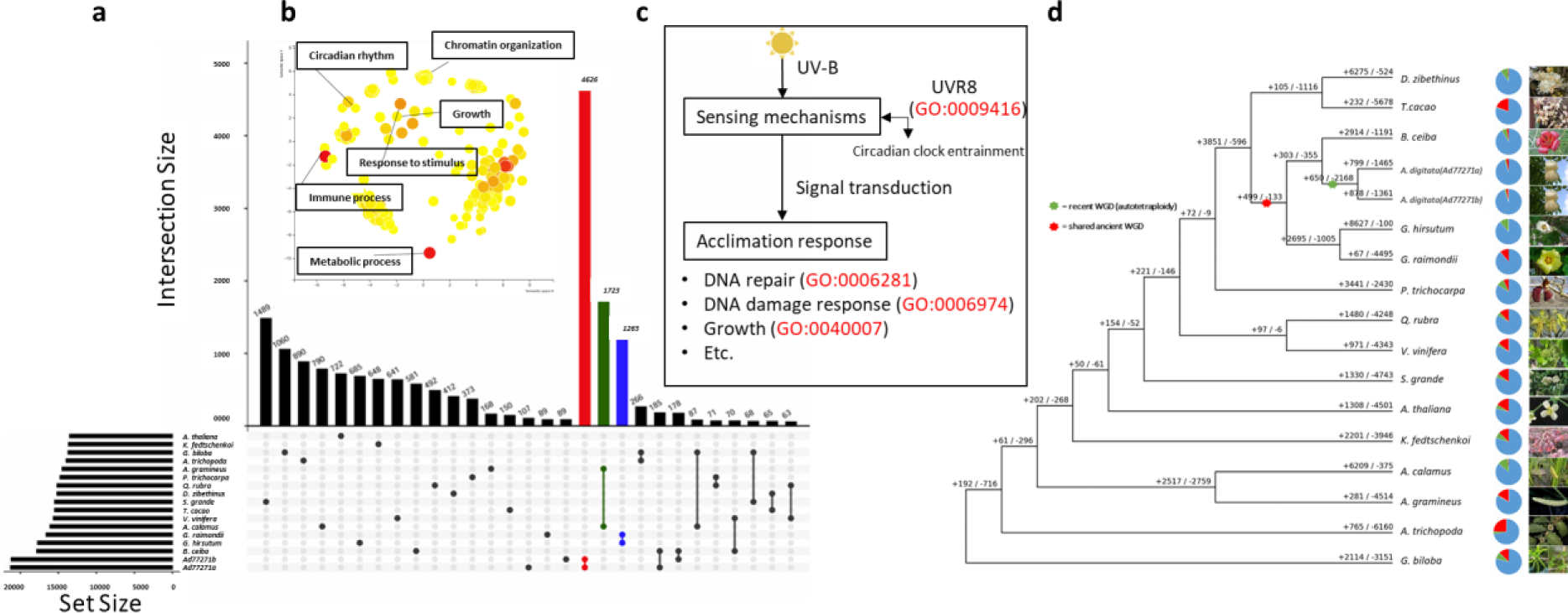
Orthogroups, gene ontology, longevity pathway, and evolutionary dynamics of African baobab tree. **a** An UpSet plot shows elevated species-specific orthogroups; red, green and blue bars correspond to baobabs, acorus (monocots), and cottons, respectively. **b** Enriched gene ontology terms in baobab include chromatin organization and circadian rhythms. **c** Ultraviolet B (UV-B) reception pathway via *UV RESISTANCE LOCUS 8* (*UVR8*) followed by clock gene entrainment, signal transduction, and acclimation responses ^49^. **d** Comparative gene evolution in baobab alongside 15 other plant species is shown in the phylogenetic tree. Positive and negative numbers represent gene family expansion and contraction, respectively. The pie chart’s blue color indicates no evident change, while green and red denote expansions and contractions in gene families. Among the compared species, diploid *Gossypium raimondii* exhibited the fewest expanded gene families. Colored asterisks represent the point of whole genome duplication, while images on the right show the plant’s inflorescence.

Leveraging gene ontology (GO) enrichment analysis, we asked which biological categories were specific to baobab. We identified 490 significant (FDR < 0.01) GO terms that could be clustered into three broad categories of metabolic processes (GO:0008152; right top), response to stimulus (GO:0050896; left top), and more disbursed group that included growth (GO:0040007), chromatin (GO:0006325) immune system process (GO:0002673) and circadian rhythm (GO:007623) (Fig. 4b; Supplementary Table 7; Supplementary Fig. 10a).

We hypothesize that this third grouping may represent genes associated with the long lived nature of baobab. For instance, there are six baobab *UV RESISTANCE LOCUS 8* (*UVR8*) genes; represented in four OG families, three of which are only found in baobab genomes. *UVR8* is a UV-A/B photoreceptor that interacts with the E3 ubiquitin-ligase *CONSTITUTIVELY PHOTOMORPHOGENIC1* (*COP1*) and *SUPPRESSOR OF PHYA-105* (*SPA*) to stabilize and destabilize two central growth-regular transcription factors *ELONGATED HYPOCOTYL 5* (*HY5*) and *PHYTOCHROME INTERACTING FACTOR 5* (*PIF5*) respectively ^49–52^ (Fig. 4c). UV-B radiation has the potential to damage macromolecules such as DNA and impair cellular processes, suggesting the baobab-specific UVR8 proteins may play a broad signaling role to protect the genome of this long lived tree. It has been hypothesized that UVR8 may interact directly with chromatin based on its orthology to *REGULATOR OF CHROMATIN CONDENSATION 1* (*RCC1*) and nucleosome binding assays ^53^. Consistent with this we observe enriched OG for DNA repair (GO:0006281), DNA damage response (GO:0006974), chromatin organization (GO:0006325) and remodeling (GO:0006338), suggesting that baobab may actively protect its genome through UVR8-chromatin regulation (Fig. 4c).

### Evidence of an ancient WGD event in baobab genome

We performed comparative genomics of the baobab genomes Ad77271a, Ad77271b, AdKB, AdOHT and Aza135 with three closely related Malvaceae species, cotton (*G. raimondii*), bombax (*B. ceiba*) and cacao (*T. cacao*), as well as grape (*V. vinifera*), which only has one whole genome triplication (WGT) and amborella (*A. trichopoda*), which is sister to the eudicot lineage and lacks a whole genome duplicates (WGD) event ^54,55^. Synteny-based and rates of synonymous substitutions (Ks) were used to estimate WGD/WGT, as well as to understand its relationship with other species, such as *A.za* (Aza135). The Ks revealed a consistent timing of the separation of baobab-amborella and baobab-grape at 128 and 96 MYA respectively (Fig. 5b) ^54,55^. Both cacao and cotton diverged from the ancestor of *Adansonia* around 30 MYA, which was the first piece of evidence that a baobab-specific WGD occurred around this time. In contrast, the bombax and Aza135 genomes diverged from Ad77271a ∼20 MYA and ∼17 MYA respectively. In addition, Aza135 had a second peak around 30 MYA suggesting it had both remnants of the WGD as well as another event separating the *Adansonia* species ^56^.

**Fig. 5:**
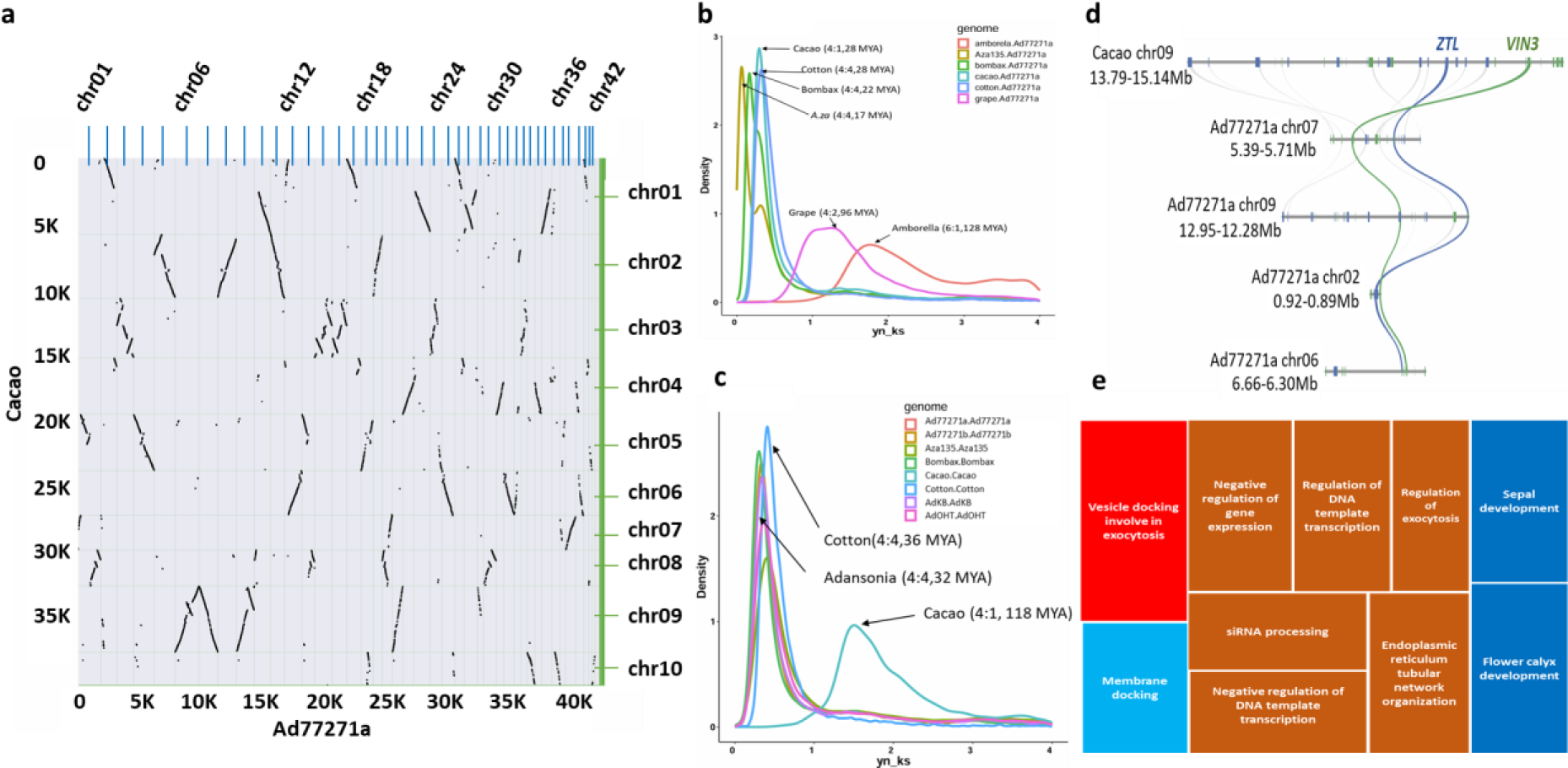
Syntenic relationships and the Ks distribution suggest an ancient whole genome duplication (WGD) event in baobab. **a** Dot plot between baobab and cacao revealing one syntenic block in the *T. cacao* genome corresponding to four distinctive syntenic blocks in the *A. digitata* genome for each *T. cacao* block, each dot represents a collinear gene pair. **b** Multi-species pairwise comparison of substitution per synonymous site (Ks); **c** Comparing Ks values within the same genome. **d** Microsynteny plot between *A. digitata* and *T. cacao* showing chromosomal region in the *T. cacao* chromosome 9 bearing gene *ZEITLUPE* (*ZTL*) and *VERNALIZATION 3* (*VIN3*) genes, which can be tracked to four regions in *A. digitata* (blue and green lines, respectively). The WGD of baobab hints unreduced polyploidy, followed by autotetraploidy. **e** Enriched Gene Ontology (GO) terms related to gene regulation for exocytosis and flower development in *A.digitata*. The syntenic depth ratio between genomes and the evolution event age in million years ago (MYA) are indicated inside the parenthesis.

The self-alignments of genomes informed the timing of the most recent WGD in a species, which clarified that all of the baobab, bombax and cotton genomes shared the same WGD event at about 30 MYA ^57^, while cacao experienced its last WGD around 118 MYA that was consistent with it having only the WGT-γ (Fig. 5c) ^58^. When we zoom into just the baobab complex, we saw that all of the genomes have a minor ks peak around 30 MYA in addition to peaks at 4, 6, 11 and 17 MYA for Ad77271b, AdKB, AdOHT and Aza135 respectively (Supplementary Fig. 7). We hypothesize that the autotetraploidy event might be seen as the distance between the Ad77271a and Ad77271b siblings since they most likely represent distinct and random haplotypes, which would place the autotetraploidy event in the 4 MYA range with a similar timing of the last detectable TEs burst (Fig. 2b).

The self-self and pairwise alignment of the haploid baobab genomes revealed a 4:4 syntenic depth consistent with a WGD event shared across *Adansonia* that has not been reduced/fractionated (Fig.s 5a; d; Supplementary Fig. 9). Cotton and bombax also shared a 4:4 syntenic depth consistent with them sharing the WGD with baobab, while cacao had a 4:1 syntenic depth, which was consistent with it not sharing the baobab WGD (Fig.s 5a; b). The 4:1 relationship between Ad77271a and cacao highlighted that the baobab has retained almost complete copies of all four chromosomes after WGD. In contrast, grape and amborella had 6:1 and 4:2 syntenic depths with Ad77271a, respectively; additionally, the baobab Ks peak left of grape in the plot is consistent with baobab having the WGT-γ and a baobab-specific WGD (Fig.s 4d; Fig. 5b).

After a WGD event, plants generally return to diploidy over a period of time and in general during this process many genes return to one copy in a process called fractionation ^59^. While whole chromosomes appeared to be retained after the WGD event in the baobab genomes, there was also some fractionation resulting in gene copies ranging from 1 to 4 (Fig. 5a; Supplementary Fig. 4d; Supplementary Table 7). We found ∼15%, 24%, 30%, and 13% of genes were retained in 1, 2, 3, and 4 copies in the Ad77271a genome after the baobab- specific WGD, which was consistent across all of the baobab genomes. We hypothesize that the genes retained as four copies in the baobab genome may represent specific biology that was important to baobab. We conducted a GO enrichment analysis of the genes that were retained as four copies and found a highly significant (bonferroni FDR < 0.05) list of overlapping GO terms that focused on gene regulation/chromatin, exocytosis, and flower timing/development (Fig. 5e).

Across most plant genomes analyzed, circadian, light and growth related genes are reduced back to one or two copies in the genome during fractionation, presumably to ensure the correct gene dosage ^59,60^. However, we found that among circadian, light and growth orthologs ^61^, only six (out of 34) genes were in one copy, while more than half (18/35) were retained as three or four copies; one pair of genes retained in four copies were an evolutionarily conserved syntenic gene pair *ZEITLUPE* (*ZTL*) and *VERNALIZATION 3* (*VIN3*), which are involved with flowering time, thermomorphogenesis, photomorphogenesis and the circadian clock, and are linked in the genome across the eudicot lineage back to amborella (Fig. 5d) ^60^. Plants that leverage Crassulacean Acid Metabolism (CAM) photosynthesis to ensure stomata are open at the correct time of day (TOD) ^62–65^, as well as the crop soybean that highly specific latitude maturity groups ^66^, have retained multiple copies of circadian, light and growth ^60^; these results could point to baobab leveraging CAM photosynthesis or some other highly regulated TOD process light flowering. These genes may provide insight into how baobab has adjusted to different environments across Africa (see variation analysis below).

### Diversity in baobab across Africa

*A. digitata* is endemic to Africa and little is known about the genetic diversity across the continent. We resequenced 25 *A. digitata* trees from across Africa using Illumina short reads, yielding an average sequencing depth of 20x per accession (Supplementary Table 1). Mapping these sequences to Ad77271a reference produced 58.9 million SNPs and 446 thousand INDELs, accessible in the <u>Baobab database (salk.edu)</u>. Analysis of ploidy using these genotypes confirmed that *A. digitata* is an autotetraploid (Supplementary Table 5). Principal component analysis (PCA) of 25 accessions revealed that more than 30% of the genetic variance was due to geographical origins, mostly east and north/west (Fig. 6a; Supplementary Table 5). Population 1 encompasses germplasms from north of the equator up to approximately 16 degrees north (and mostly west), while populations 2 and 3 span from the equator to 26 degrees south. The trees from the northern desert region of Namibia (population 3; top left; green circles) were distinct from the trees that were collected closer to Botswana (population 2). These results indicate there are geographical or environmental factors that have limited gene flow between these populations.

We leveraged the fixation index (Fst) across the three populations to determine which were more related to one another, revealing that while population 1 and 3 were more related (lower Fst; average <0.1), populations 2 and 3 as well as 1 and 3 were less related (higher average Fst >0.4) (Fig. 6b). These results were consistent with Namibian population 3 being closer to population 1, in contrast to population 2, despite their relative geographic closeness. We looked at the genes with high Fst SNPs (>0.8) across the three population comparisons to identify genes that are under selection for their environment, and only found enriched GO terms between populations 2v3 and 1v3 that could be summarized into pollination, organelle localization and chromatin (Supplementary Tables 10; 11). Consistent with our findings that baobab specific genes were related to longevity through the UVR8- chromatin connection, these genes also played a role in an environment specific fashion.

**Fig. 6:**
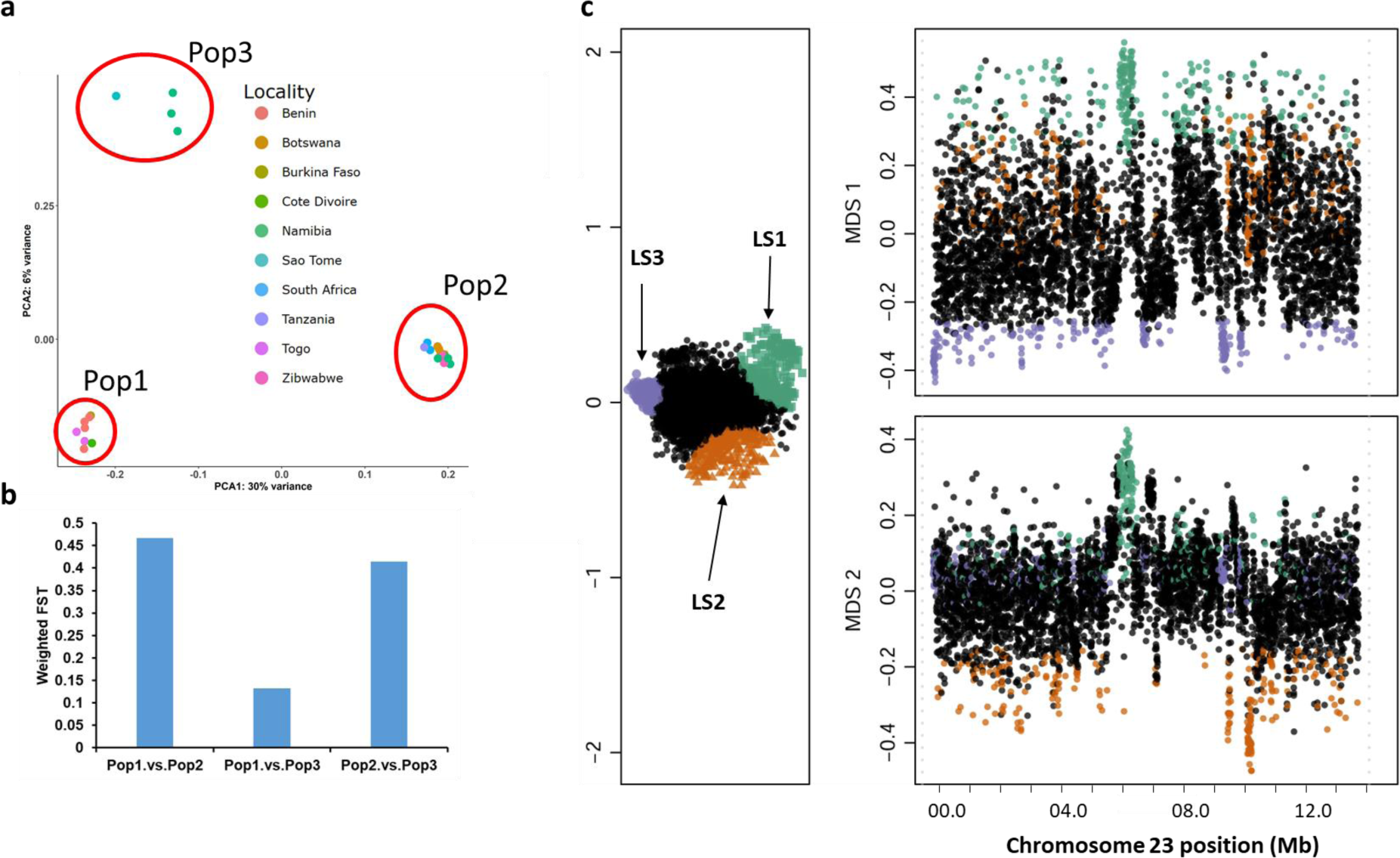
Diversity and structural variations in the *Adansonia digitata*. **a** Principal Component Analysis (PCA) of 25 *A. digitata* accessions collected across Africa using 6490 SNPs. Three clusters are present: Cluster 1 (Pop1: n=8), Cluster 2 (Pop2: n=13), Cluster 3 (Pop3: n=4). **b** Weighted FST (fixation index) values between populations: Pop1 vs. Pop2 (FST=0.47), Pop1 vs. Pop3 (FST=0.13), and Pop2 vs. Pop3 (FST=0.41). The high FST values support population differentiation due to genetic structure, primarily associated with longitudinal variation (east -west). **c** Lostruct partitions vary across chromosomes. The multidimensional scaling shows the coordinates of the 25 *A. digitata* populations on the x- axis (MDS 1) and the y-axis (MDS 2). Points that are not outliers are shown in black, while points that are outliers are shown in green (LS1), orange (LS2), and purple (LS3). Each point on the graph corresponds to a specific genomic window of 3kb. MDS stands for multidimensional scaling.

The identification of pollination genes having high Fst suggests there may be some selective pressure on synchronizing flowering and pollination, which is thought to be one of the factors that led to the ‘rise of angiosperms’ ^67,68^. Similar to CAM plants that have expanded circadian clock genes to ensure TOD specific stomatal opening, it has been hypothesized that coordination of flower development and opening must be highly TOD specific to pollinators ^60^. This may explain why many circadian clock genes have been retained in three or four copies after fractionation (Fig. 5d), and we find that many of these genes have high Fst. For instance, the circadian linked gene gene pair *ZTL* and *VIN3* that were retained in four copies also had high Fst (Fig 5d; Supplementary Table 9). Baobab flower opening is tightly regulated and only opens for one night; in East and West Africa it has been shown that bats are the primary pollinator, in southern Africa hawkmoths also play a role in pollination ^69,70^. Taken together, regulation of chromatin and pollination play important roles in shaping the diverse populations of baobab in Africa.

Variation and selection can be specific to regions of the chromosome; therefore, we employed a local PCA technique to identify patterns of relatedness from SNP frequencies across the genome ^71^. Along the chromosomes, we observed striking regions of shared relatedness that contrasted greatly from surrounding regions. One region on chromosome 23 between 6-7 Mb overlaps with the predicted centromere arrays (6-9 Mb) (Fig. 1e; 6c; Supplementary Fig.12). This region is present in Ad77271a, Ad77271b, and AdKB (all East Africa), but lost in AdOHT (Northwest Namibia), suggesting that region could be important for the diversification of baobab across Africa (Fig. 1e; Supplementary Fig.4). Many of the putative centromere regions in *A. digitata* had translocation, inversions, and duplications, or regions that didn’t map (Supplementary Fig.4; Supplementary Table 8), as well as high Fst GO enrichment of chromatin, consistent with these regions being under active change in the baobab genomes. Baobab has a large number (2*n* = 4*x* = 168) of small chromosomes (9 to 23 Mb), suggesting that selection on chromatin stability and arrangement may play a role in longevity and environmental specificity. Taken together, the baobab genome has adapted to specific environments across Africa and achieved extreme longevity by actively protecting and regulating its chromosomes, while retaining accurate TOD acuity through the retention of circadian, light and flowering time genes to ensure environment-specific pollination.

## Discussion

Baobab, an iconic succulent tree emblematic of Africa’s savannas, bears nutritious fruits fueling its global demand and bolstering income for rural communities across the continent. However, some of the oldest known baobabs are dying across Africa ^9,72^ which makes it imperative that we better understand the baobab genome to enhance yield and stress resilience. We present a high-quality chromosome-level haploid assembly of *Adansonia digitata* (Ad77271a) alongside draft genomes from a sibling tree (Ad77271b), two geographically diverse *A. digitata* (AdOHT and AdKB) and distinct species *A. za* (Aza135) from Madagascar. We also resequenced all eight *Adansonia* species and 25 *A. digitata* trees collected across Africa to estimate genome size, confirm the ploidy and estimate genetic variation. We found that the *A. digitata* genome underwent a whole genome duplication (WGD) event 4 million years ago (MYA) that resulted in autotetraploidy, which coincided with an unprecedented amplification of DNA transposable elements (TEs) compared to Long Terminal Repeat Retrotransposons (LTR-RT). In addition, baobab shares a WGD 30 MYA with the Malvaceae that displayed biased fractionation, resulting in the retention of almost full chromosomes, and multiple copies of the circadian, light and flowering pathway that could be playing a role in protecting the genome integrity for longevity as well as ensuring the timing of pollination. This research not only unravels baobab genomic evolution mysteries but also provides a crucial sequence for expediting gene discovery, enhancing breeding efforts, and aiding baobab species conservation.

While we were only able to assemble a chromosome-resolved haploid version of the *A. digitata* genome, we confirmed that it is an autotetraploid with the sibling genome (Ad77271b), centromere, WGD and the 25 resequenced trees analyses (Fig. 1d; Fig 3; Supplemental Figure 5; Supplementary Table 5). Initially, the K-mer profile of one peak with an estimated haploid genome size of 750 Mb, which is in line with what was found through Feulgen staining and flow cytometry ^1^, suggested it was diploid. However, since chromosome counting suggested that it is tetraploid ^1^, we looked more closely at the allele frequencies. Leveraging both the Ad77271b as well the resequenced 25 tree reads we analyzed coverage at heterozygous sites, which provided evidence that *A. digitata* possesses an autotetraploid genome. In addition, we identified only one centromeric repeat (base unit = 158 bp) supporting autotetraploidy, since allotetraploids like *Eragrostis tef* have been shown to have distinct centromere arrays representing the different subgenome origins ^42^. Based on the comparison with the sibling genome, the subgenomes in *A. digitata* are highly similar consistent with autopolyploids arising from the duplication of the same species’ genome(s) and thus containing four nearly identical sets of chromosomes that form tetravalents during meiosis ^73^. Finally, our Ks analysis found a WGD 4 MYA consistent with an autotetraploidy event. However, it is possible that autotetraploidy didn’t immediately lead to disomic inheritance, yet instead there may have been a long period of tetrasomic inheritances that could still be happening now as has been seen in the coast redwoods ^39^.

The baobab genome is unlike any published plant genome to date in that DNA TEs are dominant (3x) compared to LTR-RT that typically result in the bloating of plant genomes ^41^. Compared to other plant genomes, baobab has an average total TE content with *A. digitata* (∼45%) having more than *A. za* (35%), which was also in line with the 50-60% DNA methylation levels (Fig. 3; Table 2; Supplementary Fig. 8). However, the *A.digitata* TE composition was unusual compared to other plant genomes in that the proportion of DNA TEs was 33% while the LTR-RTs were 10% (Table 2). Typically, LTR-RTs are the predominant transposon in a plant genome since they proliferate in a “copy and paste” mechanism, while in contrast DNA TEs accumulate through a “cut and paste” mechanism ^41^. In terms of LTR-RT TEs, a big rise in Copia and CACTA TEs was seen around 10 MYA, which sets baobabs apart from its relative in the Malvaceae family, cacao ^55,58^. In contrast the DNA TEs burst around 4 MYA coinciding with the putative tetraploidy event in baobab, suggesting that the tetraploidy event may have played a role in the accumulation of the cut and paste DNA TEs that have shaped the baobab genome.

In addition to the WGD 4 MYA, baobab also experienced a WGD 30 MYA that is shared across the Malvaceae ^74,75^; yet unlike other Malvaceae species studied to date, baobab retained almost all four copies resulting from the WGD. While most genomes fractionate gene copies back to a diploid state after WGD, it is thought that some genes may not be fractionated and retained to modify gene dosage in specific pathways important to the organism ^59,76^. Near-complete baobab chromosomes have been retained in four copies compared to cacao (Fig. 5a; Supplementary Fig. 9), and these genes are enriched in regulation/chromatin, immunity, exocytosis, and flower development gene ontology (GO) terms (Supplemental Fig. 10). Specifically, circadian, light and flowering time genes, which are usually fractionated back to a single copy to conserve gene dosage, have been retained in multiple copies such as the evolutionarily conserved linkages between *LHY*/*PRR9* (3 copies) and *ZTL*/*VIN3* (4 copies) (Fig. 5d; Supplementary Table 9) ^60^. A similar circadian, light and flowering time gene retention was also observed in the globally important crop soybean (*Glycine max*) ^60,77^; it is thought that the extra copies may provide additional environmental specificity in soybean their internal circadian timing is correlated with their maturity groups ^66^.

The retention of circadian, light and flowering time genes provides clues as to the amazing longevity and highly regulated pollination schedule of baobab. Coupled to the observation that the baobab specific orthogroups (OG) were enriched with similar terms such as immunity, flower development, and gene regulation with a specific (Supplementary Table 7), as well as three OG with baobab-specific copies of *UV RESISTANCE LOCUS 8* (*UVR8*) (Fig. 4; Supplementary Fig. 10), led us to speculate that baobab’s longevity may be related to retention of genes in the circadian, light and flowering time pathways. Upon UV-B absorption, *UVR8* undergoes monomerization, leading to a structural change that initiates downstream signaling events through the circadian clock and *CONSTITUTIVELY PHOTOMORPHOGENIC1* (*COP1*) to regulate pathways such as DNA damage response and repair ^49,78^. In addition, genes with a high fixation index (Fst) also shared enriched terms in flower development as well as pollination. Baobab flowers only open at night and only for one night ^69,70^, suggesting that the timing of flower development and opening have to be tightly controlled for a specific environment and pollinator schedule ^60,79–82^.

Finally, we resequenced 25 *A. digitata* trees to assess the potential diversity across Africa. *Adansonia* trees situated north of the equator exhibited some divergence from those located southward, with approximately 6% of the variation attributed to this distinction. However, the most partitioning, about 30% of the variation, occurred along an east-west axis as exemplified by the distinctiveness between populations 1 and 2 (Fig. 6; Supplementary Fig. 11). Intriguingly, Namibian baobab trees clustered into two populations (2 and 3), which correspond to different watersheds, suggesting limited gene flow due to possible geographic barriers. Another study conducted in Niger and Mali supported this notion, indicating variations in baobab species across the continent; specifically, it found that West African germplasms exhibited faster growth and better adaptation to arid environments compared to their East African counterparts ^83^. Structural variation (SV) and local PCA analysis revealed that the centromere regions were highly dynamic and location specific (Fig.s 1; 6; Supplemental Fig. 4), consistent with observation that baobab may highly regulate these regions for both longevity and acuity to the specific environment.

In summary, this work presents the first chromosome-level assembly of baobab and confirms *A. digitata* as an autotetraploid species (2n=4x=168) with 42 chromosomes. WGD led to the expansion of key genes, such as circadian, light and flowering-related genes, shaping adaptation strategies for longevity and pollination in baobab ^60^. The genomic resources produced in this study will facilitate baobab genetics, conservation, and modern breeding implementation ^56,84,85^.

## Methods

### Plant material

Seeds were obtained from the USDA Germplasm Information Resource Network (GRIN) from three trees grown in USDA-Agriculture Research Service, Subtropical Horticulture Research Station, Miami, FL USA under the accession number PI77271. The original seed came from Dar es Salaam, Tanganyika Territory, Tanzania, Africa in 1928. The PI77271 tree was chosen to generate the reference genome to enable broad access to the baobab germplasm through GRIN. Seed for the *Adansonia* species to estimate genome sizes was ordered from Le Jardin Naturel (https://www.baobabs.com/). N.K. and E.H.E.K., (co-authors) collected leaf and seed material for the resequencing of 25 baobab species.

Seed was cleaned, soaked in boiling water for three days, and then planted in well drained soil. Seedlings were grown to the first true leaf stage and then dark adapted for two days for DNA and RNA extraction (Supplementary Fig. 1a).

### DNA and RNA extraction

Baobab has been a difficult species to extract high molecular weight (HMW) DNA from due to the large amounts of polysaccharides that it produces.

Therefore, we employed two different methods to obtain HMW DNA from baobab: first, seedlings at the two true leaf stages were used for DNA extraction, and second they were dark adapted for two days to deplete the polysaccharides. After two days of dark adaptation, two PI77271 seedlings were chosen for genome sequencing and named "Ad77271a" and "Ad77271b;" These sibling seedlings were chosen for sequencing to enable analysis of reported autotetraploidy. HMW DNA was extracted from Ad77271a and Ad77271b, as well as "AdOHT" from Namibia and "AdKB" from Sudan, along with *Adansonia za* (Aza135) from Madagascar using a modified protocol ^86^ (Supplementary Fig. 2). For the 25 *A. digitata* resequencing, DNA was extracted from dried leaf samples as previously described ^56^.

### Genome Sequencing

Unsheared HMW DNA (1.5 ug) from Ad77271a, Ad77271b, AdOHT, AdKB and Aza135 was used for ONT ligation-based libraries (SQK-LSK109). Final libraries were loaded on an ONT flowcell (v9.4.1) and run on the PromethION. Illumina 2x150 paired-end reads were also generated for genome size estimates and polishing genome sequences. Libraries were prepared from HMW DNA using Illumina NexteraXT library prep kit, and sequenced on NextSeq High Output 300 cycle, paired end 2X150 kit (Illumina, San Diego, CA). The resulting raw sequence was only trimmed for adaptors, resulting in >60x coverage.

### HiC library preparation

Hi-C library was prepared for "Ad77271a" using Phase Proximo HiC (Plant) kit (V.3.0) and run on Illumina NextSeq P3 300 cycle, paired end 2X150 kit.

### Genome assembly analysis

ONT reads were assembled using Flye (v2.9.2) ^87^ then polished using Racon (v1.5.0) ^88^ and Pilon (v1.24) ^89^ with Illumina reads. Hi-C data and Juicer version 1.6.2, 3ddna (v180419), and JBAT (v1.11.08) built the final assembly. The completeness of the genome assembly was assessed through BUSCO (v. 5.4.3), utilizing the ODB10 eudicots dataset ^90^.

### K-mer based genome size estimates

Applying K-mer-based techniques to Illumina short reads from Illumina sequencing libraries allowed us to estimate the genome’s size, repeat, and heterozygosity. Jellyfish in combination with GenomeScope2 ^91^ were employed to assess parameters such as haploid genome length, repeat content, and heterozygosity. For analysis of ploidy, nQuire Tool ^92^ in conjunction with statistics of variants from Illumina short reads were utilized (Supplementary Table 5).

### Scaffolding long read assembly contigs

Ragtag (v2.1.0) was used to scaffold the contigs of Ad77271b, AdOHT, AdKB and Aza135 with the HiC scaffolded Ad77271a.

### Repeats and gene prediction

EDTA (v1.9.6) ^93^ was used to construct a repeat library and softmask complex repeats. Tandem Repeats Finder (v4.09) ^94^ was employed to identify centromere and telomere sequences, as well as mask simple repeats. Genes in baobab were predicted via the Funannotate (v1.8.2) pipeline with modifications https://github.com/nextgenusfs/funannotate. Predicted proteins were characterized using Eggnog-mapper v2.0.1 ^95^.

### Long terminal repeat (LTR) insertion date

A substitution rate of 4.72 × 10−9 per year was used ^96^.

### DNA methylation analysis

LoReMe (Long Read Methylation) (<u>Oxford Nanopore</u> <u>Technologies · GitHub</u>) was used to infer DNA methylation patterns. In brief, the process involved the conversion of ONT FAST5 data to POD5 format using the Loreme Dorado- convert tool v.0.3.1. Subsequently, super-high-accuracy base calling was performed, aligning the sequences to the reference genome of *Adansonia digitata* (Ad77271a). Modkit v0.1.11 was employed to generate a bed file containing comprehensive methylation data, enabling us to create visual representations of methylation profiles for further investigation and interpretation.

### Orthology analysis

OrthoFinder (v2.5.5) ^97^ was used for comparative genomics of Malvaceae: *Adansonia digitata* (Ad77271a and Ad77271b), cotton (*Gossypium raimondii* and *Gossypium hirsutum*), cacao (*Theobroma cacao*), durio (*Durio Zibethinus*) and cotton tree (*Bombax ceiba*). Additionally, we examined representatives from Vitaceae (*Vitis vinifera*), Brassicaceae (*Arabidopsis thaliana*) Salicaceae (*Populus trichocarpa*), Fagaceae (*Quercus rubra*), Myrtaceae (*Syzygium grande*), Crassulaceae (*Kalanchoe fedtschenkoi*), Acoraceae (tetraploid *Acorus calamus* and diploid *Acorus gramineus*), Amborellaceae (*Amborella trichopoda*) and Ginkgoaceae (*Ginkgo biloba*). Except for the baobab, the primary proteins were downloaded from phytozome (v13) ^98^ and websites (Supplementary Table 6). Gene family size in the context of phylogeny was analyzed using CAFE v5.0 ^99^. Orthogroups with lots of genes in one or more species (100 genes) and only present in one species were excluded ^100^. Results were then visualized using CafePlotter (https://github.com/moshi4/CafePlotter).

### Synteny analysis

CoGe was used to make syntenic region dot plots for intergenomic and intragenomic alignments (Haug-Baltzell et al., 2017). MCscan https://github.com/tanghaibao/jcvi/wiki/MCscan-(Python-version)) was used for interspecies syntenic analysis, enabling the identification of homologous gene pairs, gene blocks, and the creation of syntenic plots that depict the relationships between homologous gene pairs between baobab and other species.

### Structural variation and rearrangement identification

Structural variations (SVs) were profiled using Syri version 1.6.3 ^101^.

### Variants analysis

In order to compare the sibling baobabs, we performed short Variant Calling and structural variant calling using two distinct pipelines: a short read (Illumina) based small variant calling workflow, and a long read (ONT) based structural variant calling workflow. Short reads were used for small variant calling due to their high accuracy at the nucleotide level, permitting high confidence SNP and short indel calls. Long reads were used for structural variant calling because their increased length allows for covering large structural variants and verifying their structure. For each of our sibling baobabs, we ran both variant calling pipelines using both baobabs as a method of sanity checking our results.

Results were consistently symmetric regardless of which of the two baobabs was used as a reference.

Our short read based short variant calling pipeline involves four primary steps: read trimming, read alignment, variant calling, quality filtering. Read trimming is helpful for filtering out low quality portions of reads that could reduce variant calling accuracy and run time as erroneous base calls when aligned to the reference have to be processed as potential variants. Read trimming was conducted using “Trim Galore” (v0.6.6). Read alignment was conducted using “minimap2” (v2.20) with short read appropriate settings and were then sorted using “Samtools” (v1.12). Each sample was mapped independently and then processed by “freebayes” (v1.3.5) collectively using tetraploid settings in order to call variants. Subsequent variant calls were then filtered to Q20 before being manually inspected and summarized with several stats tools: “vcftools” (v0.1.16) and “rtg tools” (v3.12). We also used an inhouse developed stats tool available from https://gitlab.com/NolanHartwick/bio_utils which includes functionality to process coverage stats as output by freebayes in order to verify ploidy.

### Ultraviolet-B radiation (UV-B) photoreceptor (UVR8) gene analysis

Baobab-specific orthologs were subjected to gene ontology enrichment (GO) analysis using python GOATOOLS ^102^, subsequently, visualized using REVIGO ^103^. The phylogenetic tree was analyzed via GeomeNet (https://www.genome.jp/en/about.html).

## Acknowledgements

We would like to thank the Michael Group for comments on the genome work and manuscript. We would also like to thank the genome sequencing team at Monsanto for the initial genome size survey of the *Adansonia* species funded by the Illumina Greater Good program awarded to TPM. We would also like to thank Mike Winterstein, USDA, GRIN for sending seed and leaf material from *A. digitata* tree PI77271 for genome sequencing. This work was supported by a Global Challenges Research Fund (GCRF), Nottingham Interdisciplinary Research Award and the European Research Council (ERC) under the European Union’s Horizon 2020 research and innovation programme [grant number ERC-StG 679056 HOTSPOT], via a grant to L.Y. The 25 samples used for resequencing were prepared with financial support from the National Science Foundation award DEB-1354268 to N.K. and field collecting from Diana Mayne, Sarah M. Venter, and Achille E. Assogbadjo. Finally, we are very grateful to David Baum for his constructive suggestions during the writing of the manuscript.

## Author contributions

T.P.M. and L.Y. designed the research; J.K.K., T.P.M., K.C., B.W.A., N.T.H., S.P., and N.K. performed research or analyzed data; E.H.E.K. and N.K. contributed materials and/or tools; J.K.K. and T.P.M. wrote the manuscript. All authors revised the manuscript.

## Data availability

The complete Ad77271a, Ad77271b, AdOHT, AdKB and Aza135 genomes are available through the Michael Lab genome portal: Baobab database (salk.edu), and are also uploaded to CoGe ID: 67790, 67791, 67792, 67793, 67794, 67795, 67796, 67797,67798, 67799,

67800, and 67801. In addition, the genomes and raw data can be accessed under BioProject 1022505: http://www.ncbi.nlm.nih.gov/bioproject/1022505.

---

**Supplementary Fig. 1:**
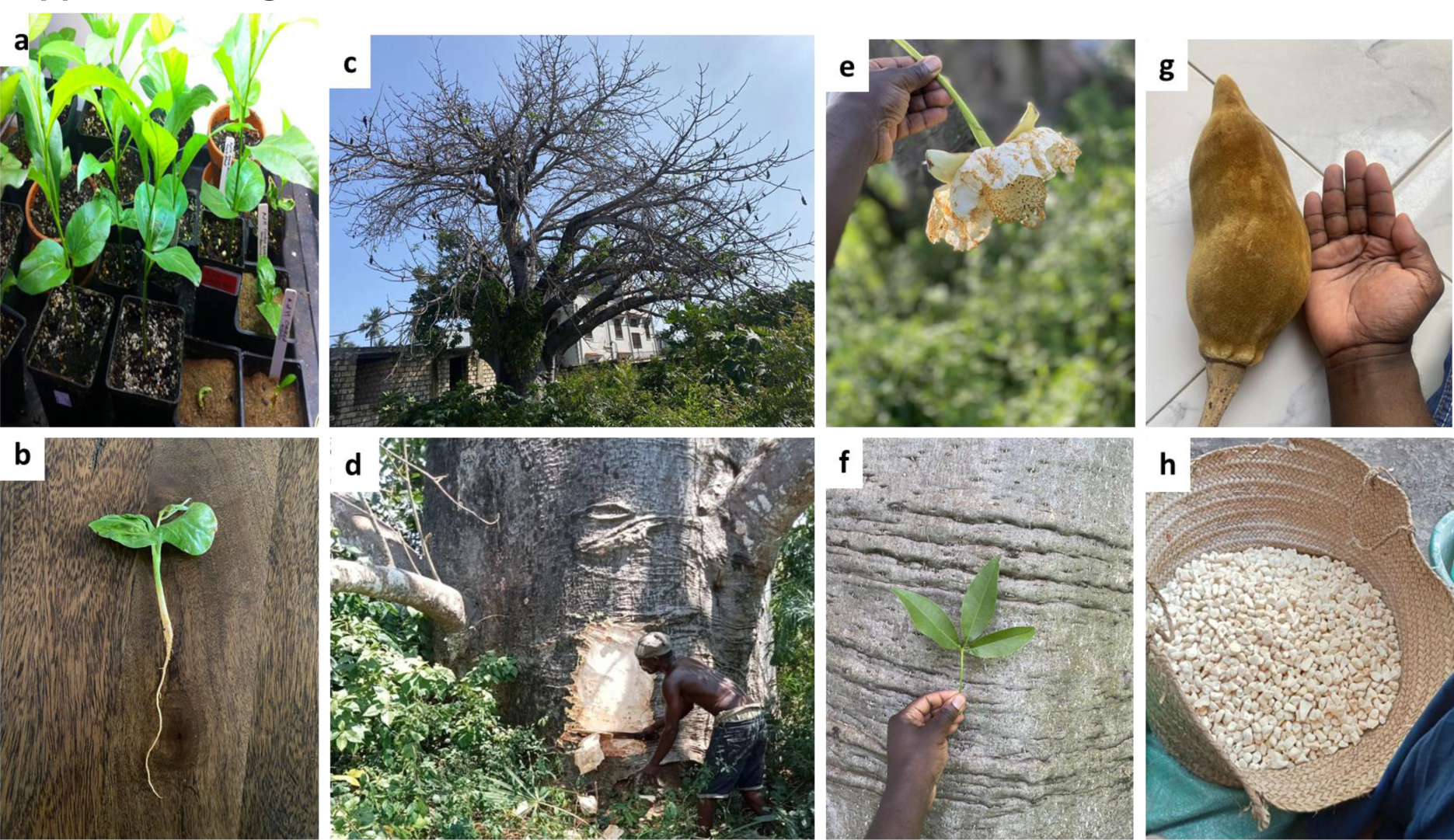
Baobab (*Adansonia digitata*) is an economically important tree. **a** USDA-GRIN PI 77271 seedlings used for genome sequencing. **b** seedling tap root. **c** Deciduous mature tree in its natural habitat: a representation of a long history of coexistence with humans. **d** Bark harvesting for fiber production in Kwale, Kenya. **e** Whitish waxy flower that has a diameter of upto 20cm (8"). **f** Shiny reflective grayish surface of mature bark and juvenile leaf. **g** Yellowish hard woody pod of mature fruit with lengths of upto 30cm (12"). **h** Powdery, whitish fruit pulp, which is abundant in vitamin C, antioxidants, calcium, potassium, and dietary fiber.

**Supplementary Fig. 2:**
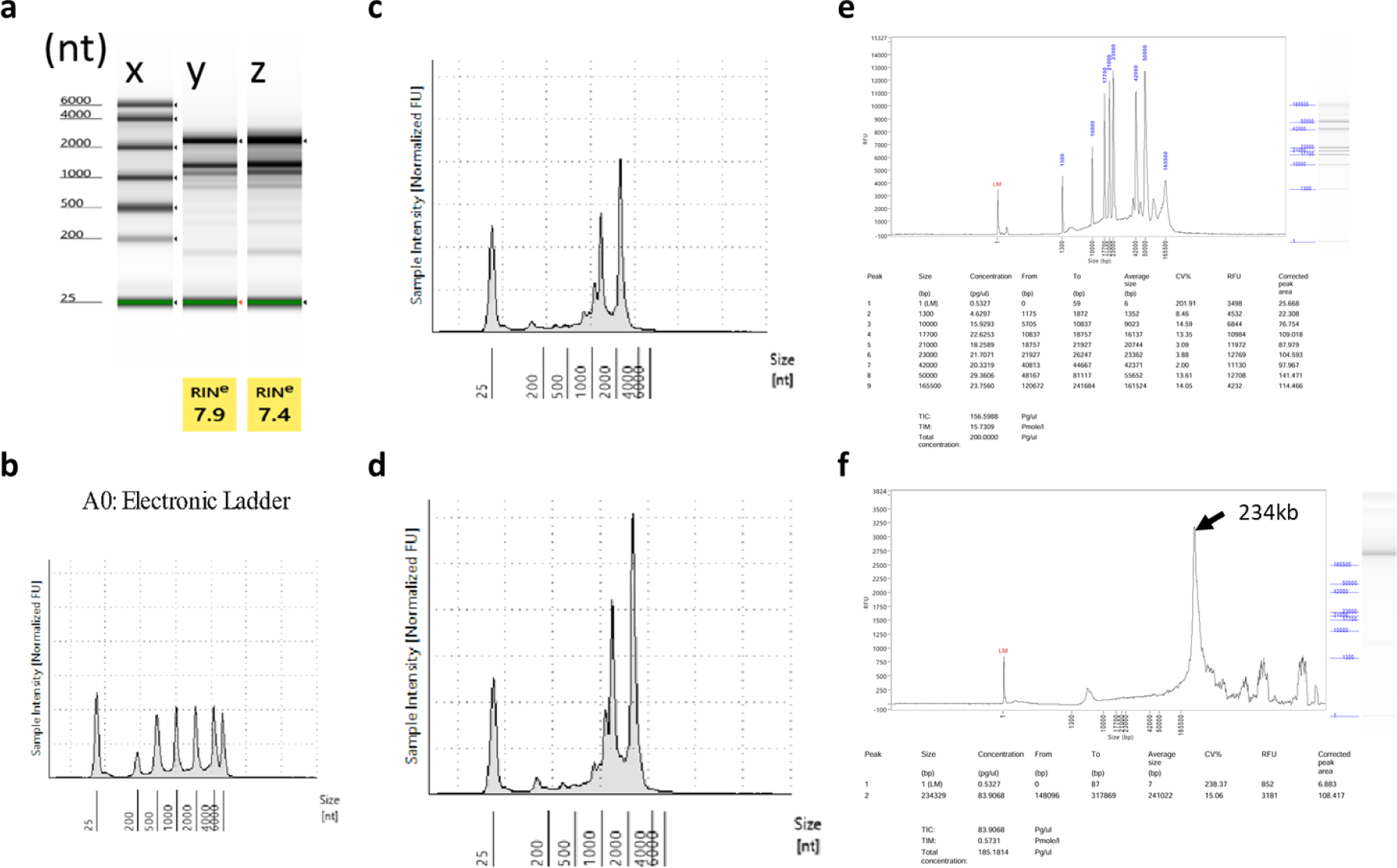
RNA and DNA quality summary. **a** Gel image and RNA Integrity Number (RIN) values for Ad77271a and Ad77271b RNA; x corresponds to the electronic ladder, y is Ad77271a, and z is Ad77271b. **b** RNA electronic ladder. **c** Electropherograms for Ad77271a . **d** Electropherograms for Ad77271b. **e** DNA electronic ladder. **f** Baobab DNA run on Femto pulse, majority of the DNA was over 165kb in length with main peak estimated at 234kb.

**Supplementary Fig. 3:**
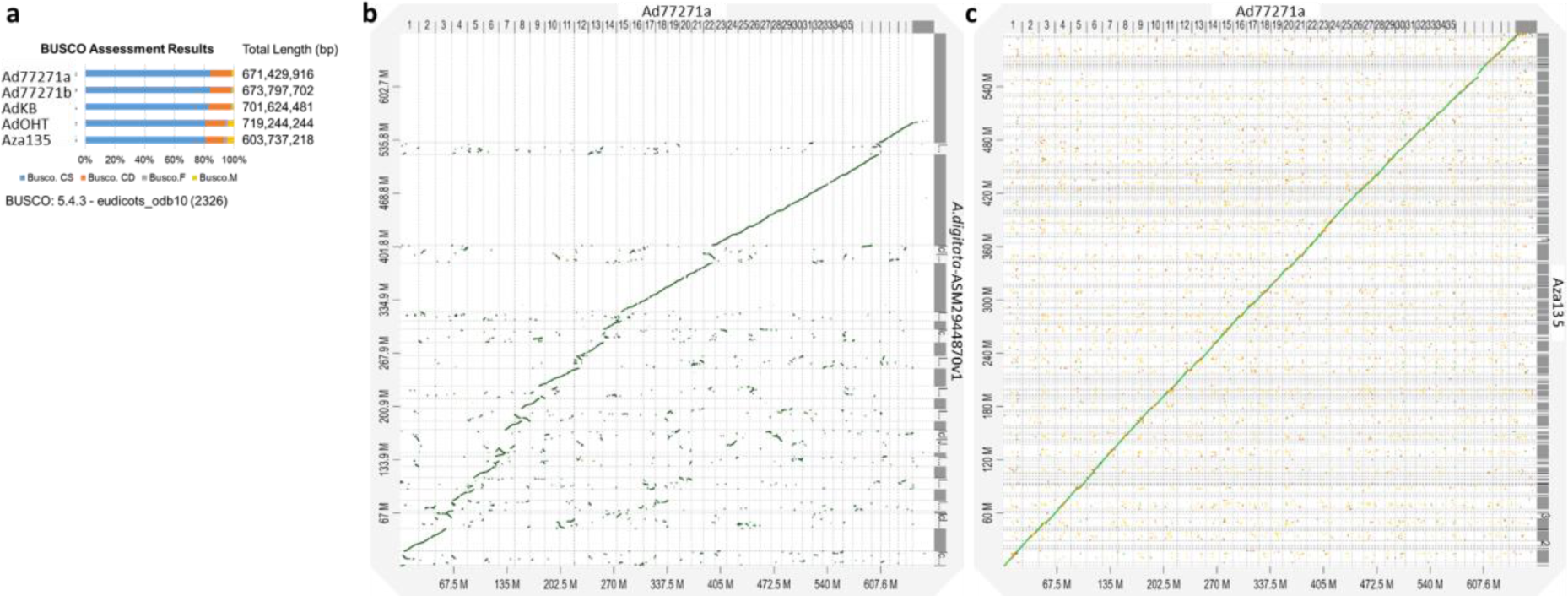
Evaluation of baobab genome assemblies. **a** Benchmarking Universal Single-Copy Orthologs (BUSCO) evaluation of Ad77271a, Ad77271b, AdKB, AdOHT and Aza135 baobab genomes. **b** Syntenic plot comparing sequences of Ad77271a against *Adansonia digitata*-ASM2944870v1 (woods et al., 2023) genomes. **c** Synteny dot plot of Ad77271a against Aza135 genome.

**Supplementary Fig. 4:**
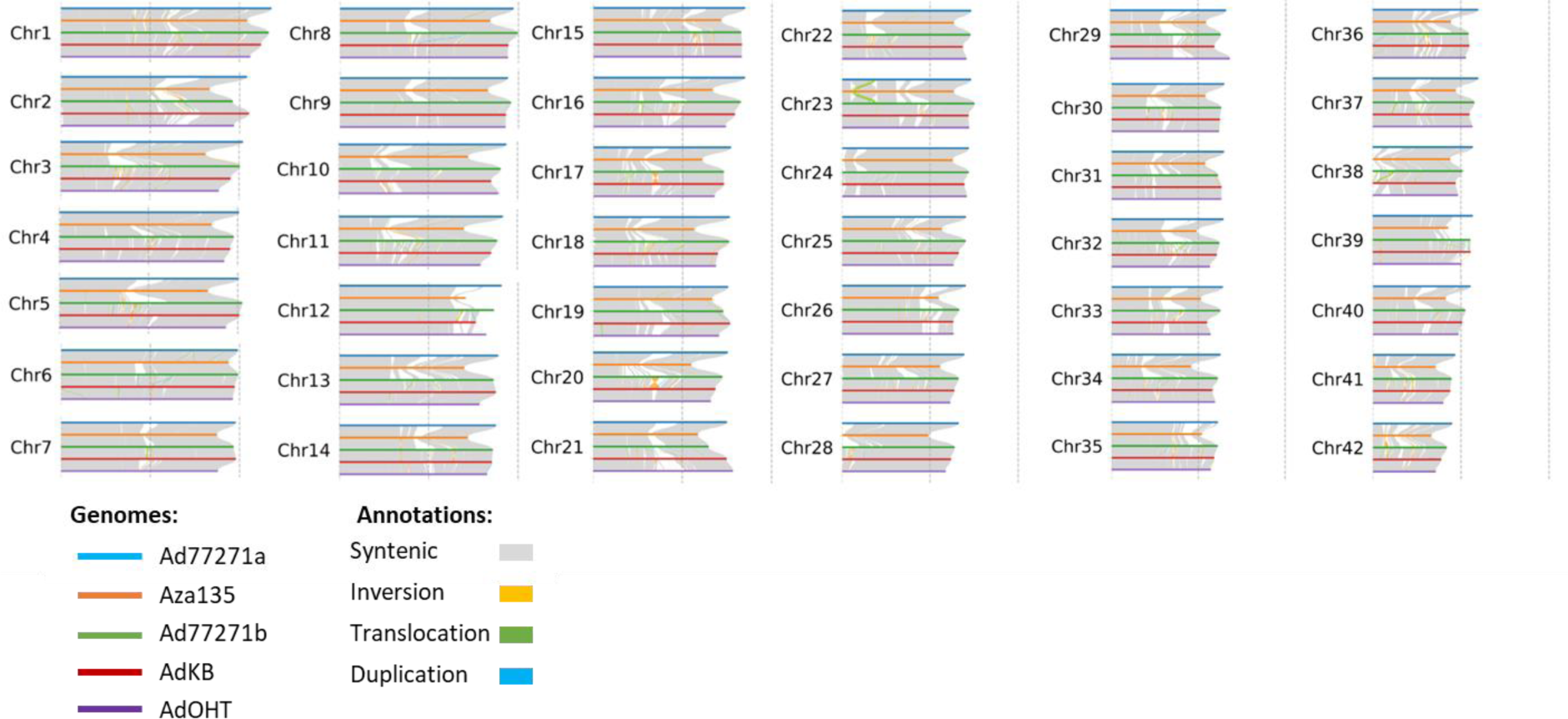
Structural rearrangements and synteny between *A. digitata* (Ad77271a, Ad77271b, AdKB, and AdOHT) and *A. za* (Aza135). Gray, orange, green, and blue ribbon colors represent syntenic, inversion, translocation, and duplication structural variations, respectively.

**Supplementary Fig. 5:**
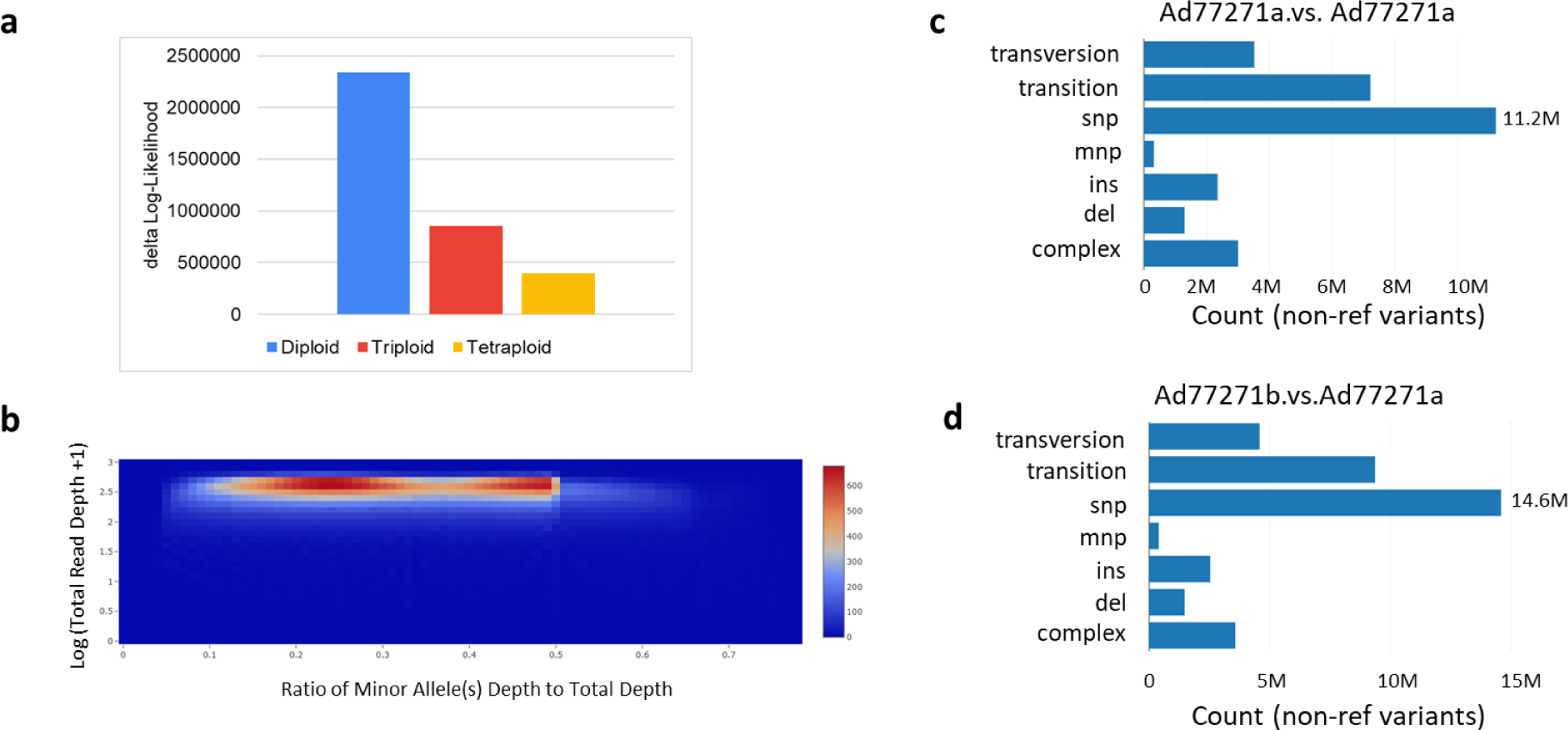
Variant analysis reveals autotetraploidy in the *Adansonia digitata* genome. **a** Gaussian Mixture Model (GMM) estimation of ploidy. This models frequency distributions at variant sites with two segregating bases and uses maximum likelihood to pick the most likely model. The ploidy level with the smallest Δ logL is identified as the true ploidy (tetraploid for baobab). **b** Two-dimensional histogram illustrates ploidy based on minor allele frequency coverage for Ad77271b; the sibling genome of Ad77271a. For diploid organisms, a single peak is expected. However, for tetraploid organisms, the histogram should exhibit two peaks, approximately located at 0.25 and 0.5. The summary of different variants is shown for **c** Ad77271a vs. Ad77271a **d** Ad77271b vs. Ad77271a. For diploid loci and homozygous alternates, it would contribute two points. For heterozygous, it would contribute one point.

**Supplementary Fig. 6:**
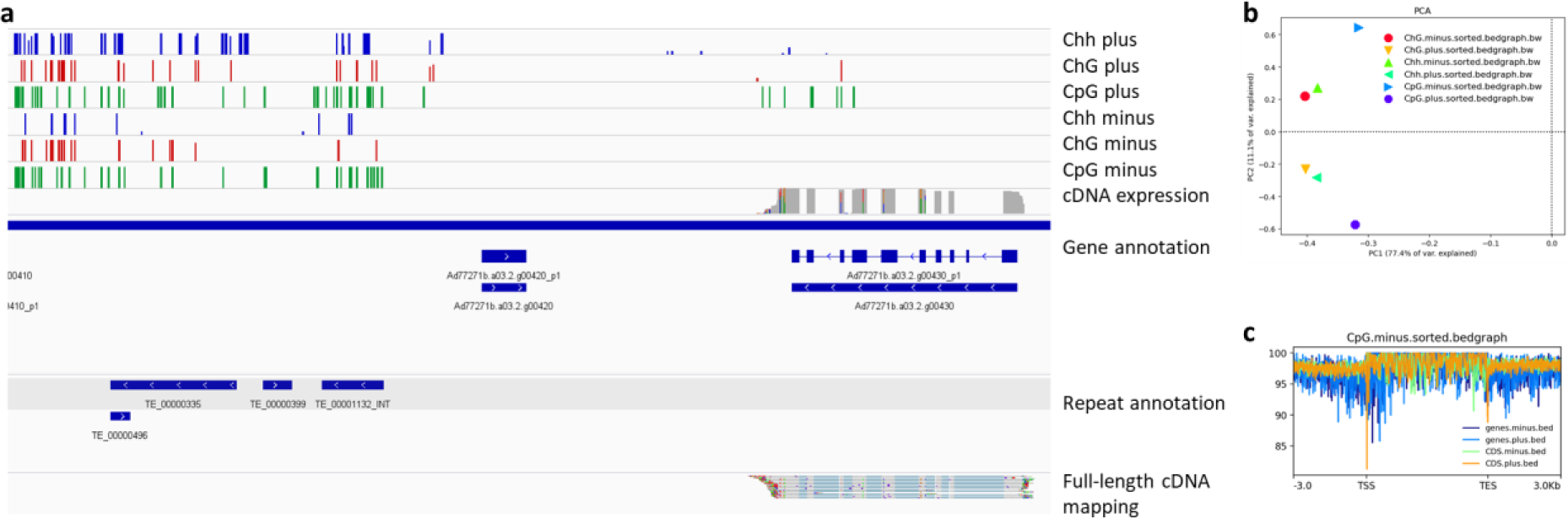
Unveiling methylation patterns in baobab genome. **a** Hypermethylation in transposable elements contrasted with hypomethylation of genes. **b** Correlation of methylation on the same strand with varied 5mC methylation types; and **c** Enhanced methylation in gene bodies and specific coding regions compared to intergenic regions.

**Supplementary Fig. 7:**
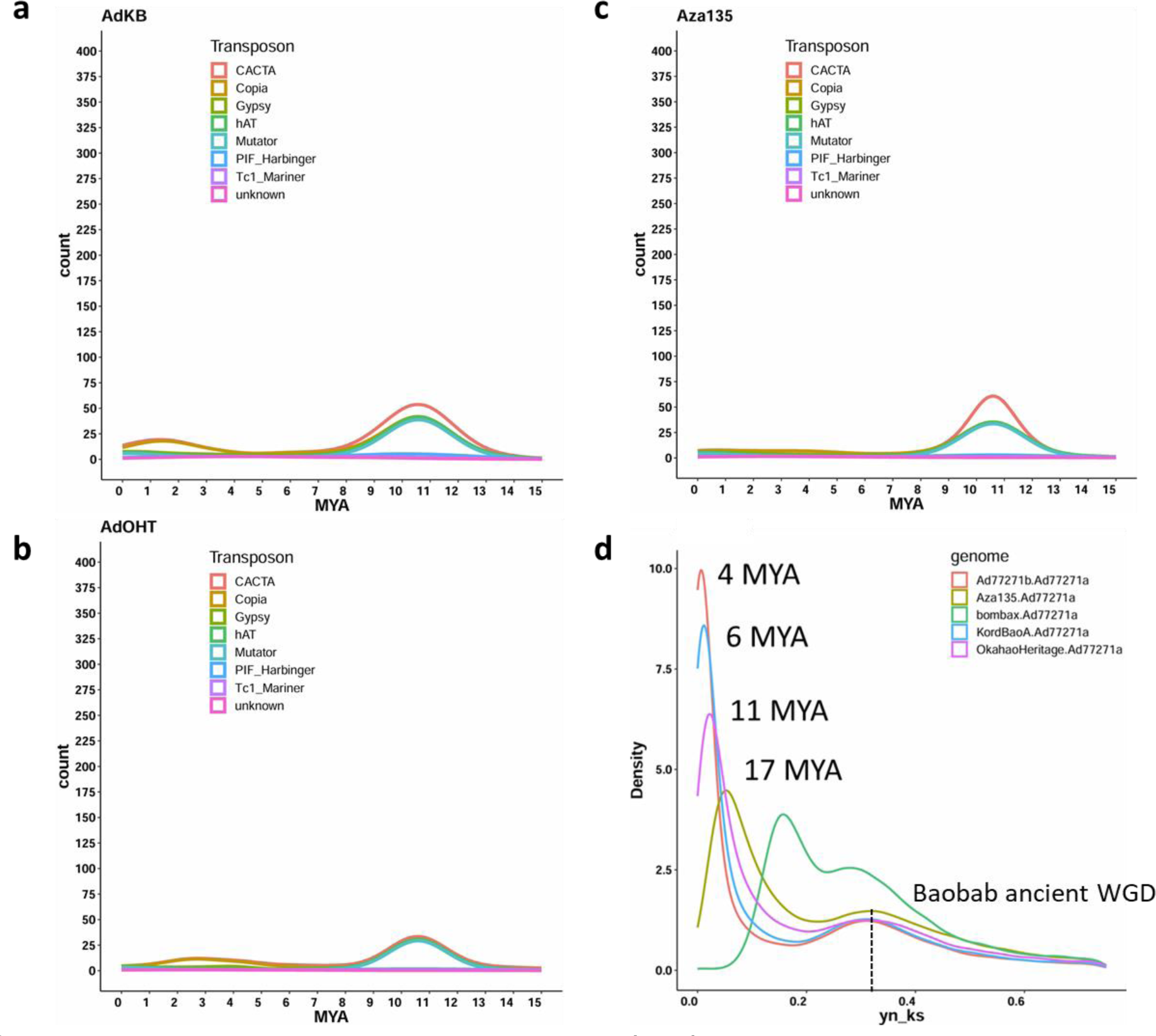
Transposable elements (TEs) evolution and baobab whole genome duplication (WGD) events. **a**, **b** and **c** Density plots for AdKB, AdOHT and Aza135 showing intact TEs burst relative to estimated insertion periods in Million Years Ago (MYA). Around 10-12 MYA, baobab genome experienced elevated levels of CACTA and COPIA long terminal repeat retrotransposon(LTR-RTs). Additionally, 3-4 MYA TEs were proliferated. **d** ks (synonymous substitution rate) distribution in baobab. We hypothesize the distance between the Ad77271a and Ad77271b siblings to be time of autotetraploidization since they most likely represent distinct and random haplotypes. The peaks at 4, 6, 11 and 17 MYA for Ad77271b, AdKB, AdOHT and Aza135 regions show baobab accessions splits.

**Supplementary Fig. 8:**
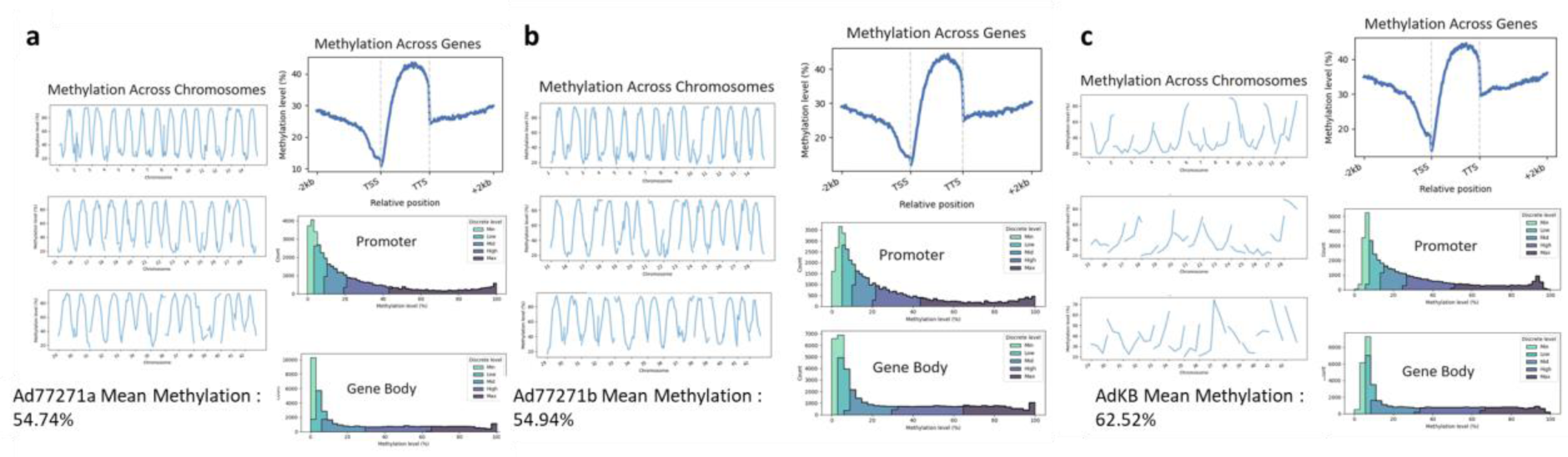
Assessment of methylation levels in baobab genomes. Analysis of methylation across the 42 chromosomes (first column) and mean percentage methylation; methylation patterns across genes, promoters, and gene bodies (second column) for genomes: **a** Ad77271a, **b** Ad77271b, and **c** AdKB.

**Supplementary Fig. 9:**
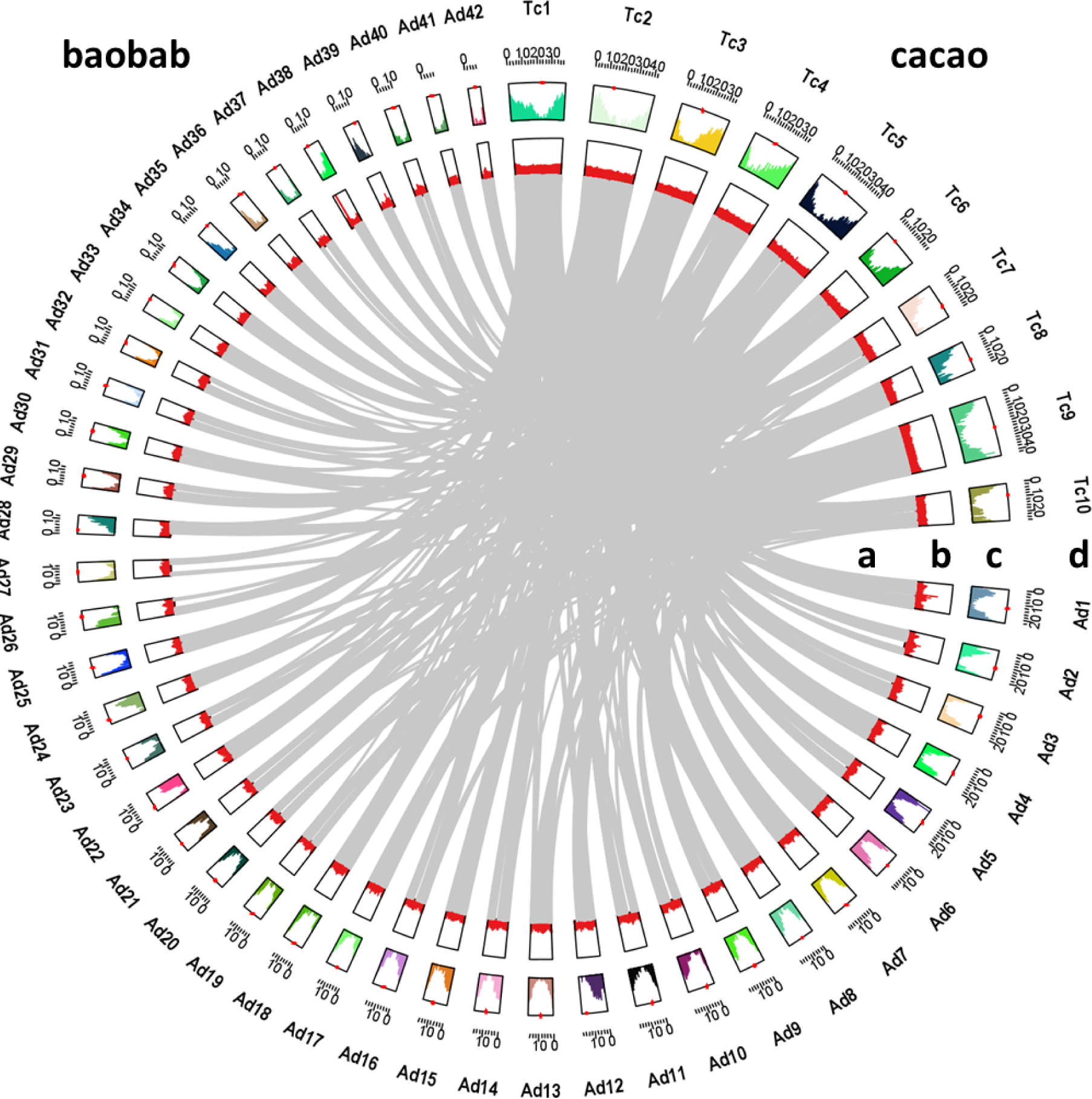
Comparative analysis of autotetraploid *A. digitata* and diploid *Theobroma cacao* genomes. Inner to outer tracks depict: a Syntenic genes, b GC content, c Gene density, and d Chromosome information. Prefixes ’Ad’ and ’Tc’ denote baobab and cacao respectively. The circos plot illustrates 42 pseudomolecules for baobab and 10 for cacao, with a window size of 100 kb. The red asterisk highlights the metacentric and acrocentric centromeres in baobab.

**Supplementary Fig. 10:**
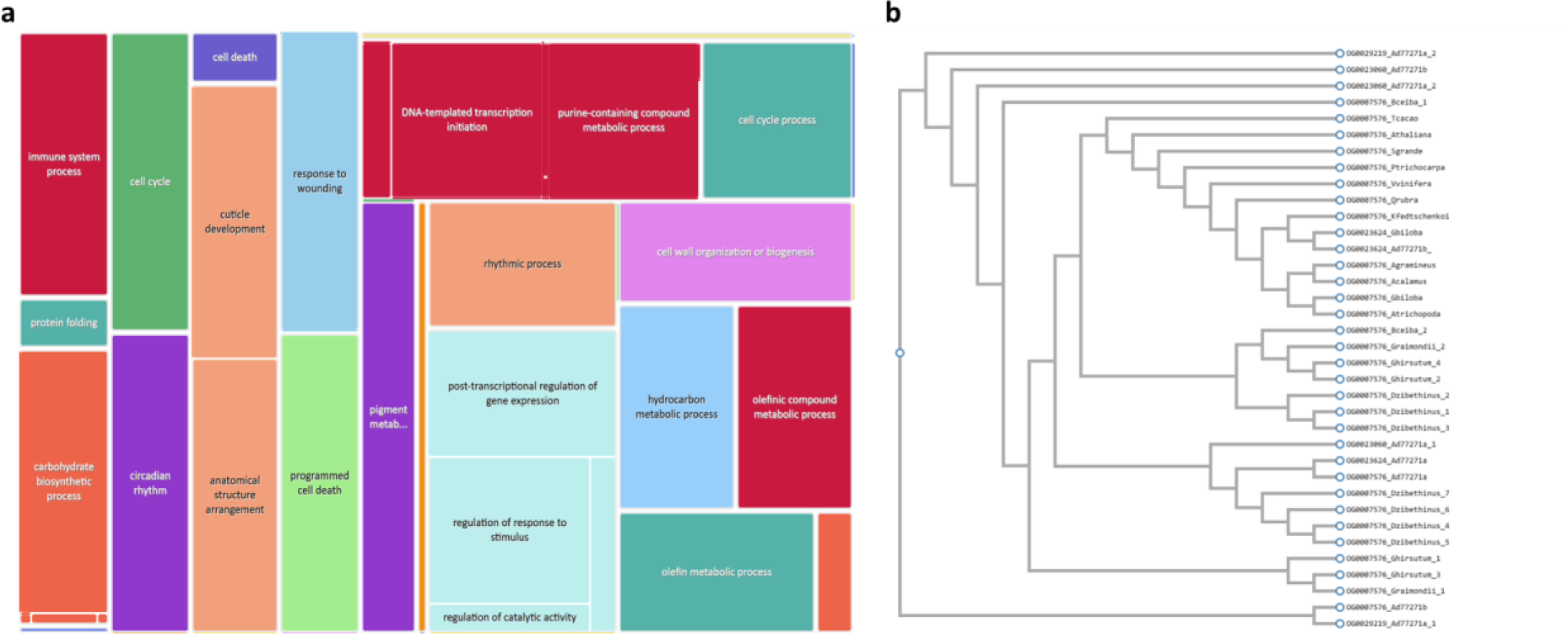
Biological process genes expanded during baobab evolution. **a** Bar chart displaying GO terms significantly enriched in *Adansonia digitata* (Ad77271a) genes. Cluster of terms related to stress response, including response to wounding (GO:0009611), cell death (GO:0008219) and circadian rhythm (GO:007623). **b** Comparative analysis reveals six ultraviolet-B receptor like genes (UVR8) in Ad77271a; the genes cluster into four orthogroups, of which three are unique to *A. digitata* (OG0029219, OG0023060 and OG0023624).

**Supplementary Fig. 11:**
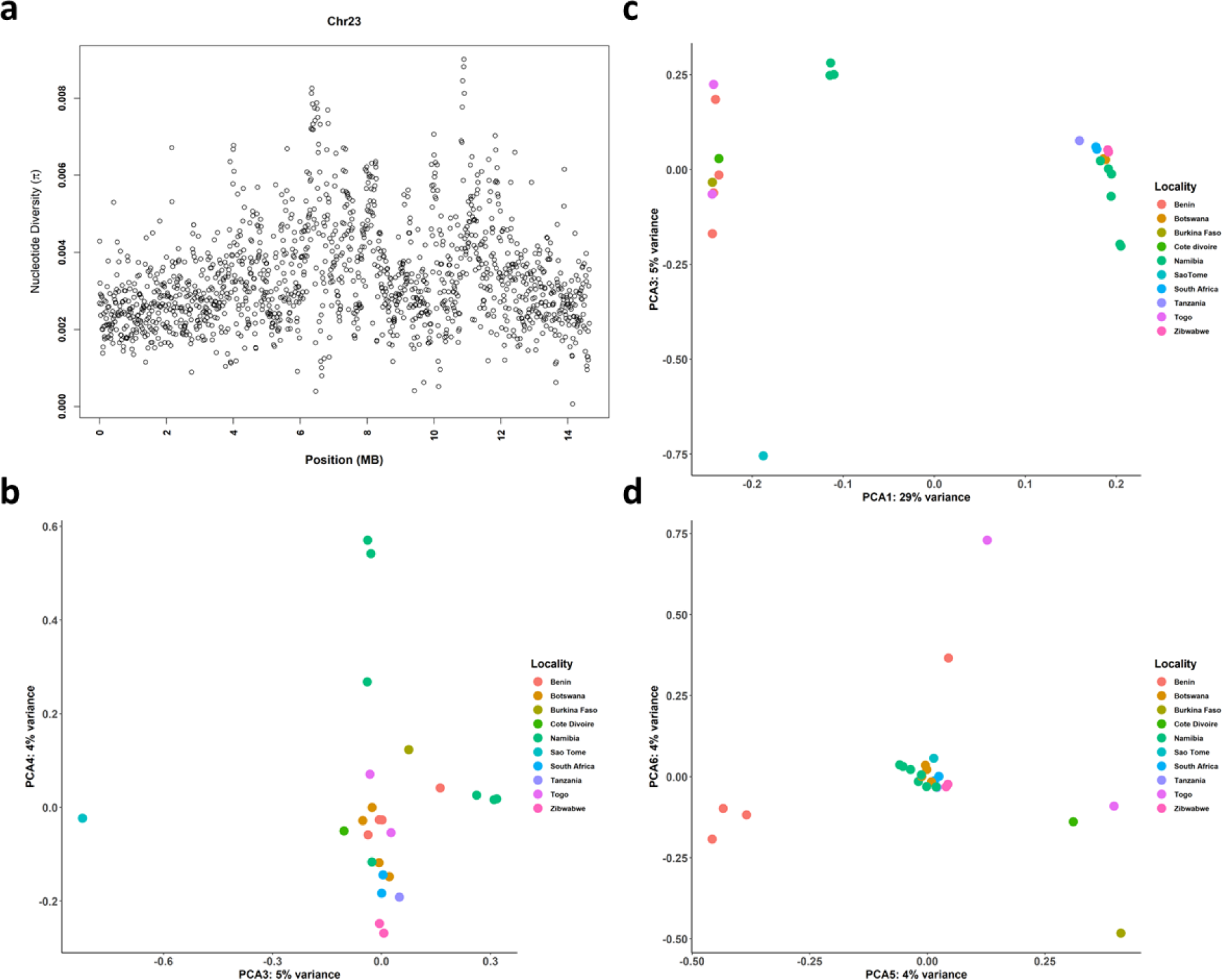
Genetic diversity and population structure of African baobab. **a** Nucleotide diversity along chromosome 23 in 25 African baobab populations. Chromosome 23 harbored a translocation distinguishing *A. digitata* from *A. za* species (Fig. 1e). Principal Component Analysis (PCA) colored by locality. Panels represent clustering using 6490 SNPs in 25 Adansonia populations: **b** axis 3 vs axis 4; **c** axis 1 vs axis 3; **d** axis 5 vs axis 6.

**Supplementary Fig. 12:**
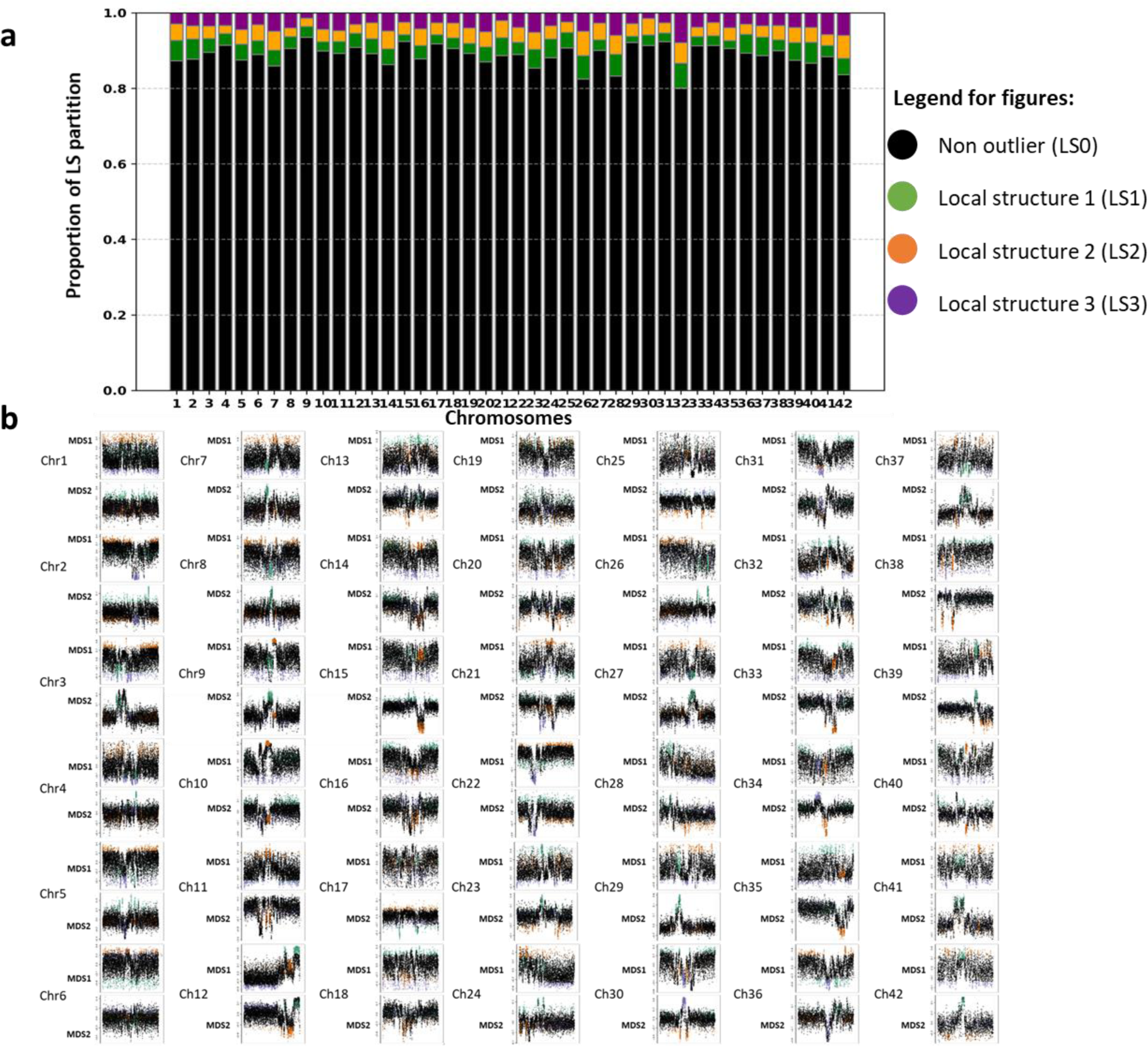
Lostruct partitions vary across *A. digitata* chromosomes. **a** Proportion of chromosomes assigned to LS0 (black), LS1 (green), LS2 (orange), LS3 (purple) in lostruct partitions. Lostruct uses principal component analysis and multidimensional scaling (MSDS) ^71^. **b** Local population structure analysis revealed outlier subsets on the 42 chromosomes. We identified three distinct outlier subsets, labeled LS1 (green), LS2 (orange), and LS3 (purple). These subsets were then compared against the rest of the genome, which represents the non-outliers (black) on 3kb windows. Chromosome sizes are not to scale.

## References

1. Islam-Faridi, N., Sakhanokho, H. F. & Dana Nelson, C. New chromosome number and cyto-molecular characterization of the African Baobab (Adansonia digitata L.) - “The Tree of Life.” Sci. Rep. 10, 13174 (2020).

2. Gibb, H. A. R. & Beckingham, C. F. The Travels of Ibn Battuta*, AD* 1325–1354. (Taylor & Francis, 1999).

3. Baum, D. A. A Systematic Revision of Adansonia (Bombacaceae). (Missouri Botanical Garden, 1995).

4. Asogwa, I. S., Ibrahim, A. N. & Agbaka, J. I. African baobab: Its role in enhancing nutrition, health, and the environment. *Trees For*. People 3, 100043 (2021).

5. Silva, V. M., Putti, F. F., White, P. J. & Reis, A. R. D. Phytic acid accumulation in plants: Biosynthesis pathway regulation and role in human diet. Plant Physiol. Biochem. 164, 132–146 (2021).

6. Rashford, J. Baobab. (Springer Cham, 2023).

7. 7. Research & Markets ltd. Baobab Powder: Global Strategic Business Report. researchandmarkets.com https://www.researchandmarkets.com/reports/5029822/baobab-powder-global-strategic-business-report (2023).

8. Offiah, V. O. & Falade, K. O. Potentials of baobab in food systems. *Appl*. Food Res 3, 100299 (2023).

9. Patrut, A. et al. The demise of the largest and oldest African baobabs. Nat Plants 4, 423–426 (2018).

10. Gebauer, J. et al. Africa’s wooden elephant: the baobab tree (Adansonia digitata L.) in Sudan and Kenya: a review. Genet. Resour. Crop Evol. 63, 377–399 (2016).

11. Venter, S. M. & Witkowski, E. T. F. Where are the young baobabs? Factors affecting regeneration of Adansonia digitata L. in a communally managed region of southern Africa. J. Arid Environ. 92, 1–13 (2013).

12. Venter, S. M. et al. Baobabs (Adansonia digitata L.) are self-incompatible and “male” trees can produce fruit if hand-pollinated. S. Afr. J. Bot. 109, 263–268 (2017).

13. Karimi, N. et al. Evidence for hawkmoth pollination in the chiropterophilous African baobab (Adansonia digitata). Biotropica 54, 113–124 (2021).

14. Harris, B. J. & Baker, H. G. Pollination of flowers by bats in Ghana. Nigerian Field 24, 151–159 (1959).

15. Jaeger, P. Epanouissement et pollinisation fleur du Baobab. Compt. Rend. Hebd. S6ances Acad. Sci. 220, 369–372 (1945).

16. Coe, M. J. & Isaac, F. M. Pollination of the baobab (Adansonia digitata L.) by the lesser bush baby (Galago crassicaudatus E. Geoffroy). East Afr. Wildl. J 3, 123–124 (1965).

17. Cron, G. V. et al. One African baobab species or two? Synonymy of Adansonia kilima and A. digitata. TAXON 65, 1037–1049 (2016).

18. Patrut, A. et al. Radiocarbon dating of two old African baobabs from India. PLoS One. 15, e0227352 (2020).

19. Kitony, J. K. Nested association mapping population in crops: current status and future prospects. J. Crop Sci. Biotechnol. 26, 1–12 (2022).

20. Wickens, G. E. The Baobabs: Pachycauls of Africa, Madagascar and Australia. (Springer Netherlands, 2008).

21. Chan, E. K. F. et al. Human origins in a southern African palaeo-wetland and first migrations. Nature 575, 185–189 (2019).

22. Sanchez, A. C., Patrick, E., Osborne & Haq, N. Climate change and the African baobab (Adansonia digitata L.): the need for better conservation strategies. Afr J Ecol 49, 234– 245 (2011).

23. Wild, S. Africa’s majestic baobab trees are mysteriously dying. Nature. 10.1038/d41586-018-05411-7<x> (2018) doi:10.1038/d41586-018-05411-7.

24. Woods, S., O’Neill, K. & Pirro, S. The Complete Genome Sequence of (Malvaceae, Malvales), the African Baobab. Biodivers Genomes 2023, (2023).

25. Costa, L., Oliveira, Á., Carvalho-Sobrinho, J. & Souza, G. Comparative cytomolecular. analyses reveal karyotype variability related to biogeographic and species richness patterns in Bombacoideae (Malvaceae). Plant Syst. Evol. 303, 1131–1144 (2017).

26. Baum, D. A. & Oginuma, K. A review of chromosome numbers in Bombacaceae with new counts for Adansonia. TAXON 43, 11–20 (1994).

27. Pettigrew FRS, J. D. et al. Morphology, ploidy and molecular phylogenetics reveal a new diploid species from Africa in the baobab genus Adansonia (Malvaceae: Bombacoideae). TAXON 61, 1240–1250 (2012).

28. Bennett, M. D. & Leitch, I. J. Nuclear DNA amounts in angiosperms: targets, trends and tomorrow. Ann. Bot. 107, 467–590 (2011).

29. Henniges, M. C. et al. The Plant DNA C-Values Database: A One-Stop Shop for Plant Genome Size Data. Methods Mol. Biol. 2703, 111–122 (2023).

30. Sun, H. et al. Chromosome-scale and haplotype-resolved genome assembly of a tetraploid potato cultivar. Nat. Genet. 54, 342–348 (2022).

31. Aklilu, B. B. et al. Functional Diversification of Replication Protein A Paralogs and Telomere Length Maintenance in Arabidopsis. Genetics 215, 989–1002 (2020).

32. Aklilu, B. B., Soderquist, R. S. & Culligan, K. M. Genetic analysis of the Replication Protein A large subunit family in Arabidopsis reveals unique and overlapping roles in DNA repair, meiosis and DNA replication. Nucleic Acids Res. 42, 3104–3118 (2014).

33. Ishibashi, T. et al. Two types of replication protein A in seed plants. FEBS J. 272, 3270– 3281 (2005).

34. Takashi, Y., Kobayashi, Y., Tanaka, K. & Tamura, K. Arabidopsis replication protein A 70a is required for DNA damage response and telomere length homeostasis. Plant Cell Physiol. 50, 1965–1976 (2009).

35. Colt, K. et al. Telomere Length in Plants Estimated with Long Read Sequencing. bioRxiv. 2024.03.27.586973 (2024) doi:10.1101/2024.03.27.586973.

36. Whittemore, K., Vera, E., Martínez-Nevado, E., Sanpera, C. & Blasco, M. A. Telomere shortening rate predicts species life span. Proc. Natl. Acad. Sci. U. S. A. 116, 15122– 15127 (2019).

37. Han, Y. et al. Chromosome-level genome assembly of Welwitschia mirabilis, a unique Namib Desert species. Mol. Ecol. Resour. 22, 391–403 (2022).

38. Wan, T. et al. The Welwitschia genome reveals a unique biology underpinning extreme longevity in deserts. Nat. Commun. 12, 4247 (2021).

39. Scott, A. D., Stenz, N. W. M., Ingvarsson, P. K. & Baum, D. A. Whole genome duplication in coast redwood (Sequoia sempervirens) and its implications for explaining the rarity of polyploidy in conifers. New Phytol. 211, 186–193 (2016).

40. Ernst, E. et al. The genomes and epigenomes of aquatic plants (Lemnaceae) promote triploid hybridization and clonal reproduction. bioRxiv 2023.08.02.551673 (2023) doi:10.1101/2023.08.02.551673.

41. Michael, T. P. Plant genome size variation: bloating and purging DNA. Brief. Funct. Genomics 13, 308–317 (2014).

42. VanBuren, R. et al. Exceptional subgenome stability and functional divergence in the allotetraploid Ethiopian cereal teff. Nat. Commun. 11, 884 (2020).

43. Bell, C. G. et al. DNA methylation aging clocks: challenges and recommendations. Genome Biol. 20, 249 (2019).

44. Wilkinson, G. S. et al. DNA methylation predicts age and provides insight into exceptional longevity of bats. Nat. Commun. 12, 1615 (2021).

45. Mira, S., Pirredda, M., Martín-Sánchez, M., Marchessi, J. E. & Martín, C. DNA methylation and integrity in aged seeds and regenerated plants. Seed Sci. Res. 30, 92– 100 (2020).

46. Gallego-Bartolomé, J. DNA methylation in plants: mechanisms and tools for targeted manipulation. New Phytol. 227, 38–44 (2020).

47. Naish, M. et al. The genetic and epigenetic landscape of the Arabidopsis centromeres. Science 374, eabi7489 (2021).

48. Niederhuth, C. E. et al. Widespread natural variation of DNA methylation within angiosperms. Genome Biol. 17, 194 (2016).

49. Tilbrook, K. et al. The UVR8 UV-B Photoreceptor: Perception, Signaling and Response. Arabidopsis Book 11, e0164 (2013).

50. Tossi, V. E. et al. Beyond Arabidopsis: Differential UV-B Response Mediated by UVR8 in Diverse Species. Front. Plant Sci. 10, 780 (2019).

51. Liu, W. et al. Phosphorylation of Arabidopsis UVR8 photoreceptor modulates protein interactions and responses to UV-B radiation. Nat. Commun. 15, 1221 (2024).

52. Jenkins, G. I. The UV-B Photoreceptor UVR8: From Structure to Physiology. Plant Cell. 26, 21–37 (2014).

53. Bourbousse, C., Barneche, F. & Laloi, C. Plant Chromatin Catches the Sun. Front. Plant Sci. 10, 1728 (2019).

54. Amborella Genome Project. The Amborella genome and the evolution of flowering plants. Science 342, 1241089 (2013).

55. Jaillon, O. et al. The grapevine genome sequence suggests ancestral hexaploidization in major angiosperm phyla. Nature 449, 463–467 (2007).

56. Karimi, N. et al. Reticulate Evolution Helps Explain Apparent Homoplasy in Floral Biology and Pollination in Baobabs (Adansonia; Bombacoideae; Malvaceae). Syst. Biol. 69, 462–478 (2020).

57. Conover, J. L. et al. A Malvaceae mystery: A mallow maelstrom of genome multiplications and maybe misleading methods? J. Integr. Plant Biol. 61, 12–31 (2019).

58. Argout, X. et al. The genome of Theobroma cacao. Nat. Genet. 43, 101–108 (2010).

59. Cheng, F. et al. Gene retention, fractionation and subgenome differences in polyploid plants. Nat Plants 4, 258–268 (2018).

60. Michael, T. P. Core circadian clock and light signaling genes brought into genetic linkage across the green lineage. Plant Physiol. 190, 1037–1056 (2022).

61. Lou, P. et al. Preferential retention of circadian clock genes during diploidization following whole genome triplication in Brassica rapa. Plant Cell 24, 2415–2426 (2012).

62. Wickell, D. et al. Underwater CAM photosynthesis elucidated by Isoetes genome. Nat. Commun. 12, 6348 (2021).

63. Yang, X. et al. The Kalanchoë genome provides insights into convergent evolution and building blocks of crassulacean acid metabolism. Nat. Commun. 8, 1899 (2017).

64. Wai, C. M. et al. Time of day and network reprogramming during drought induced CAM photosynthesis in Sedum album. PLoS Genet. 15, e1008209 (2019).

65. Ming, R. et al. The pineapple genome and the evolution of CAM photosynthesis. Nat. Genet. 47, 1435–1442 (2015).

66. Greenham, K. et al. Geographic Variation of Plant Circadian Clock Function in Natural and Agricultural Settings. J. Biol. Rhythms 32, 26–34 (2017).

67. Condamine, F. L., Silvestro, D., Koppelhus, E. B. & Antonelli, A. The rise of angiosperms pushed conifers to decline during global cooling. Proc. Natl. Acad. Sci. U. S. A. 117, 28867–28875 (2020).

68. Soltis, P. S., Folk, R. A. & Soltis, D. E. Darwin review: angiosperm phylogeny and evolutionary radiations. Proceedings of the Royal Society B: Biological Sciences 286, 20190099 (2019).

69. Chetty, A., Glennon, K. L., Venter, S. M., Cron, G. V. & Witkowski, E. T. F. Reproductive ecology of the African baobab: Floral features differ among individuals with different fruit production. For. Ecol. Manage. 489, 119077 (2021).

70. Taylor, P. J., Vise, C., Krishnamoorthy, M. A., Kingston, T. & Venter, S. Citizen Science Confirms the Rarity of Fruit Bat Pollination of Baobab (Adansonia digitata) Flowers in Southern Africa. Diversity 12, 106 (2020).

71. Li, H. & Ralph, P. Local PCA Shows How the Effect of Population Structure Differs Along the Genome. Genetics 211, 289–304 (2019).

72. Wild, S. Africa’s majestic baobab trees are mysteriously dying. Nature Preprint at 10.1038/d41586-018-05411-7 (2018).

73. Blanc, G. & Wolfe, K. H. Widespread paleopolyploidy in model plant species inferred from age distributions of duplicate genes. Plant Cell 16, 1667–1678 (2004).

74. Wang, K. et al. The draft genome of a diploid cotton Gossypium raimondii. Nat. Genet. 44, 1098–1103 (2012).

75. Kim, Y.-M. et al. Genome analysis of Hibiscus syriacus provides insights of polyploidization and indeterminate flowering in woody plants. DNA Res. 24, 71–80 (2017).

76. Feng, X. et al. Genomic evidence for rediploidization and adaptive evolution following the whole-genome triplication. Nat. Commun. 15, 1635 (2024).

77. Garsmeur, O. et al. Two evolutionarily distinct classes of paleopolyploidy. Mol. Biol. Evol. 31, 448–454 (2014).

78. Fehér, B. et al. Functional interaction of the circadian clock and UV RESISTANCE LOCUS 8-controlled UV-B signaling pathways in Arabidopsis thaliana. Plant J. 67, 37– 48 (2011).

79. Marshall, C. M., Thompson, V. L., Creux, N. M. & Harmer, S. L. The circadian clock controls temporal and spatial patterns of floral development in sunflower. Elife 12, (2023).

80. Fenske, M. P., Nguyen, L. P., Horn, E. K., Riffell, J. A. & Imaizumi, T. Circadian clocks of both plants and pollinators influence flower seeking behavior of the pollinator hawkmoth Manduca sexta. Sci. Rep. 8, 2842 (2018).

81. Bloch, G., Bar-Shai, N., Cytter, Y. & Green, R. Time is honey: circadian clocks of bees and flowers and how their interactions may influence ecological communities. Philos. Trans. R. Soc. Lond. B Biol. Sci. 372, (2017).

82. Fenske, M. P. & Imaizumi, T. Circadian Rhythms in Floral Scent Emission. Front. Plant Sci. 7, 462 (2016).

83. Korbo, A. et al. Comparison of East and West African populations of baobab (Adansonia digitata L.). Agrofor. Syst. 85, 505–518 (2011).

84. Liao, N. et al. Chromosome-level genome assembly of bunching onion illuminates genome evolution and flavor formation in Allium crops. Nat. Commun. 13, 6690 (2022).

85. Budhlakoti, N. et al. Genomic Selection: A Tool for Accelerating the Efficiency of Molecular Breeding for Development of Climate-Resilient Crops. Front. Genet. 13, 832153 (2022).

86. Lutz, K. A., Wang, W., Zdepski, A. & Michael, T. P. Isolation and analysis of high quality nuclear DNA with reduced organellar DNA for plant genome sequencing and resequencing. BMC Biotechnol. 11, 54 (2011).

87. Kolmogorov, M., Yuan, J., Lin, Y. & Pevzner, P. A. Assembly of long, error-prone reads using repeat graphs. Nat. Biotechnol. 37, 540–546 (2019).

88. Vaser, R., Sović, I., Nagarajan, N. & Šikić, M. Fast and accurate de novo genome assembly from long uncorrected reads. Genome Res. 27, 737–746 (2017).

89. Walker, B. J. et al. Pilon: an integrated tool for comprehensive microbial variant detection and genome assembly improvement. PLoS One 9, e112963 (2014).

90. Manni, M., Berkeley, M. R., Seppey, M., Simão, F. A. & Zdobnov, E. M. BUSCO Update: Novel and Streamlined Workflows along with Broader and Deeper Phylogenetic Coverage for Scoring of Eukaryotic, Prokaryotic, and Viral Genomes. Mol. Biol. Evol. 38, 4647–4654 (2021).

91. Ranallo-Benavidez, T. R., Jaron, K. S. & Schatz, M. C. GenomeScope 2.0 and Smudgeplot for reference-free profiling of polyploid genomes. Nat. Commun. 11, 1432 (2020).

92. Weiß, C. L., Pais, M., Cano, L. M., Kamoun, S. & Burbano, H. A. nQuire: a statistical framework for ploidy estimation using next generation sequencing. BMC Bioinform 19, 1–8 (2018).

93. Ou, S. et al. Author Correction: Benchmarking transposable element annotation methods for creation of a streamlined, comprehensive pipeline. Genome Biol. 23, 76 (2022).

94. Benson, G. Tandem repeats finder: a program to analyze DNA sequences. Nucleic Acids Res. 27, 573–580 (1999).

95. Cantalapiedra, C. P., Hernández-Plaza, A., Letunic, I., Bork, P. & Huerta-Cepas, J. eggNOG-mapper v2: Functional Annotation, Orthology Assignments, and Domain Prediction at the Metagenomic Scale. Mol. Biol. Evol. 38, 5825–5829 (2021).

96. Cossu, R. M., Buti, M., Giordani, T., Natali, L. & Cavallini, A. A computational study of the dynamics of LTR retrotransposons in the Populus trichocarpa genome. Tree Genet. Genomes 8, 61–75 (2011).

97. Emms, D. M. & Kelly, S. OrthoFinder: phylogenetic orthology inference for comparative genomics. Genome Biol. 20, 238 (2019).

98. Goodstein, D. M. et al. Phytozome: a comparative platform for green plant genomics. Nucleic Acids Res. 40, D1178–86 (2012).

99. Fábio K Mendes, Dan Vanderpool, Ben Fulton and Matthew W Hahn. CAFE 5 models variation in evolutionary rates among gene families. Bioinformatics 36, 5516–5518 (2020).

100. Padgitt-Cobb, L. K., Pitra, N. J., Matthews, P. D., Henning, J. A. & Hendrix, D. A. An improved assembly of the “Cascade” hop () genome uncovers signatures of molecular evolution and refines time of divergence estimates for the Cannabaceae family. Hortic Res 10, uhac281 (2023).

101. Goel, M., Sun, H., Jiao, W.-B. & Schneeberger, K. SyRI: finding genomic rearrangements and local sequence differences from whole-genome assemblies. Genome Biol. 20, 277 (2019).

102. Klopfenstein, D. V. et al. GOATOOLS: A Python library for Gene Ontology analyses. Sci. Rep. 8, 1–17 (2018).

103. Supek, F., Bošnjak, M., Škunca, N. & Šmuc, T. REVIGO summarizes and visualizes long lists of gene ontology terms. PLoS One 6, e21800 (2011).

